# GMIP-PLSR: A Nextflow Pipeline for GWAS and Multi-Omics Integration in Gene Prioritization Using PLSR

**DOI:** 10.64898/2026.04.06.716845

**Authors:** Mohammed S Kanchwala, Chao Xing, Zhenyu Xuan

## Abstract

Genome-wide association studies (GWAS) have significantly advanced our understanding of complex traits and diseases, but their interpretive power remains limited due to challenges in identifying causal genes and pathways. Integrating GWAS with multi-omics data—such as gene expression, protein-protein interactions, and gene-pathway networks have the potential to enhance biological insights and improve gene prioritization. To fulfill this potential and need, we developed the GWAS & Multi-omics Integration Pipeline (GMIP), a flexible and scalable framework that incorporates widely used tools such as PoPS, MAGMA, and benchmarker to enrich GWAS findings. However, PoPS suffers from multicollinearity in its features, which can impact performance. To overcome this, we introduce GMIP-PLSR, an extension of GMIP that uses Partial Least Squares Regression (PLSR) to manage multicollinearity effectively. We applied GMIP-PLSR across multiple GWAS datasets, demonstrating superior performance over PoPS in most cases. In a case study on NAFLD, GMIP-PLSR, using features derived from both disease-specific scRNA-seq and general PoPS features, identified gene sets with higher heritability and stronger enrichment in known NAFLD pathways, confirming its ability to enhance GWAS findings. Built on Nextflow, GMIP is computationally efficient, adaptable to diverse research environments, and provides a robust solution for gene reprioritization in post-GWAS analyses.

GMIP-PLSR is available at https://github.com/mohammedmsk/GMIP.

## 1.0 Introduction

Genetics serves as a fundamental scientific discipline, offering critical insights into the mechanisms underlying biological traits and diseases (Johnson et al., 2007). The advent of technologies like genome sequencing, bioinformatics, and high-throughput methods has profoundly expanded our understanding of complex diseases, opening new avenues for diagnosis and therapy (Lander et al., 2001). The molecular mechanisms by which genes influence biological traits are central to understanding disease, with variations in processes like transcription and translation leading to a range of disorders (Strachan & Read, 2010). Genome-wide association studies (GWAS) have accelerated the identification of genes associated with specific traits or diseases by examining genetic variants across populations, linking them to complex conditions such as diabetes, Alzheimer’s, and cardiovascular disease (Visscher et al., 2012a). GWAS plays a vital role in drug discovery by identifying novel drug targets (Plenge et al., 2013). Furthermore, pharmacogenetics, which investigates how genetic variation influences drug response, is increasingly important for developing personalized therapies that optimize treatment efficacy and minimize adverse effects based on an individual’s genetic makeup (Rodríguez-Antona & Taron, 2015).

Despite the transformative impact of GWAS, a significant challenge remains in directly identifying causal genes or pathways. Most GWAS-identified loci contain numerous genetic variants in linkage disequilibrium (LD), making it difficult to pinpoint the exact causal variant (Visscher et al., 2017; Wood et al., 2014b). Moreover, a substantial proportion of these variants are in non-coding regions of the genome, with their functional roles and target genes often remaining elusive (Maurano et al., 2012). Pinpointing causal genes is crucial for gaining fundamental insights into disease pathogenesis, developing targeted therapeutic interventions, understanding shared genetic architectures across diseases (facilitating drug repurpose), and broadly comprehending human biology and genetic variation (Hindorff et al., n.d.; Visscher et al., 2012b; Sanseau et al., 2012; Solovieff et al., 2013; Lonsdale et al., 2013; Manolio et al., 2009b)

To overcome these limitations, the integration of GWAS with multi-omics datasets has emerged as a powerful approach. Multi-omics integration combines data from various molecular layers, including genomics, transcriptomics, proteomics, and epigenomics, to provide a more comprehensive understanding of the biological mechanisms underlying complex traits (Gamazon et al., 2015; Hasin et al., 2017; Lonsdale et al., 2013; Marbach et al., 2016; Wu et al., 2019). This integration allows researchers to validate GWAS findings, uncover novel associations, and gain deeper insights into the functional impact of genetic variants. Existing integrative methods include locus-based colocalization tools like eCAVIAR (Hormozdiari et al., 2016), COLOC (G. Wang et al., 2020), ENLOC (Wen et al., 2017), and mediation or association methods like TWAS (Gusev et al., 2016), SMR (Zhu et al., 2016), and similarity-based methods like NetWAS (C. S. Greene et al., 2015), NAGA (Carlin et al., 2019), GPrior (Kolosov et al., 2021). However, these methods are often used in isolation, with limited systematic assessments of their power and error rates, and many do not utilize the full array of available multi-omics datasets (Fine et al., 2019). Recently, the Polygenic Priority Score (PoPS) pipeline has been made available, leveraging the full polygenic signal in GWAS summary statistics and incorporating gene features from a variety of sources, including 73 publicly available scRNA-seq datasets (Weeks et al., 2023). It has been shown to be better than most of the locus-based and similarity-based methods (Weeks et al., 2023).

Despite these advancements, two major challenges persist in gene prioritization:

1. **Lack of a Uniform Framework:** There is no standardized framework for comparing and integrating various gene prioritization methods. Different tools operate independently, using distinct feature sets and analytical strategies, which complicates cross-comparisons and limits the systematic evaluation and optimization of these methods. This fragmented landscape prevents researchers from exploring cross-combinations of modules or approaches, which could potentially lead to better gene prioritization results.
2. **Multicollinearity in Feature Sets:** Many gene prioritization methods suffer from multicollinearity, where features (such as expression levels and network interactions) are highly correlated. This redundancy can reduce the ability of methods like PoPS to accurately isolate causal genes, affecting the overall performance of prioritization tasks. Addressing this issue is critical to improve the precision and reliability of gene ranking, particularly in multi-omics datasets (Piccininni et al., 2022).

To address these critical gaps, we developed the **G**WAS and **M**ulti-omics **I**ntegration **P**ipeline (GMIP), a modularized Nextflow pipeline. GMIP offers a unified framework to compare and integrate various gene prioritization methods, thereby addressing the lack of standardization in the field. Furthermore, we extended GMIP to **GMIP-PLSR** by incorporating Partial Least Squares Regression (PLSR). This integration directly tackles the issue of multicollinearity, particularly in feature-rich methods like PoPS, leading to enhanced gene prioritization performance and improved result interpretability. Our results demonstrate that GMIP with PLSR consistently improves gene prioritization across various GWAS datasets, outperforming existing methods in both accuracy and interpretability.

## 2.0 Materials and Methods

### 2.1 GMIP Framework Overview

GMIP is structured around four essential modules: SNP2Gene Mapping, Machine Learning Modeling, Cross-Validation Strategy, and Evaluation. These modules are designed to facilitate gene prioritization by integrating GWAS data with multi-omics information. Figure 1 in results section illustrates the overall framework of GMIP.

**Figure 1:**
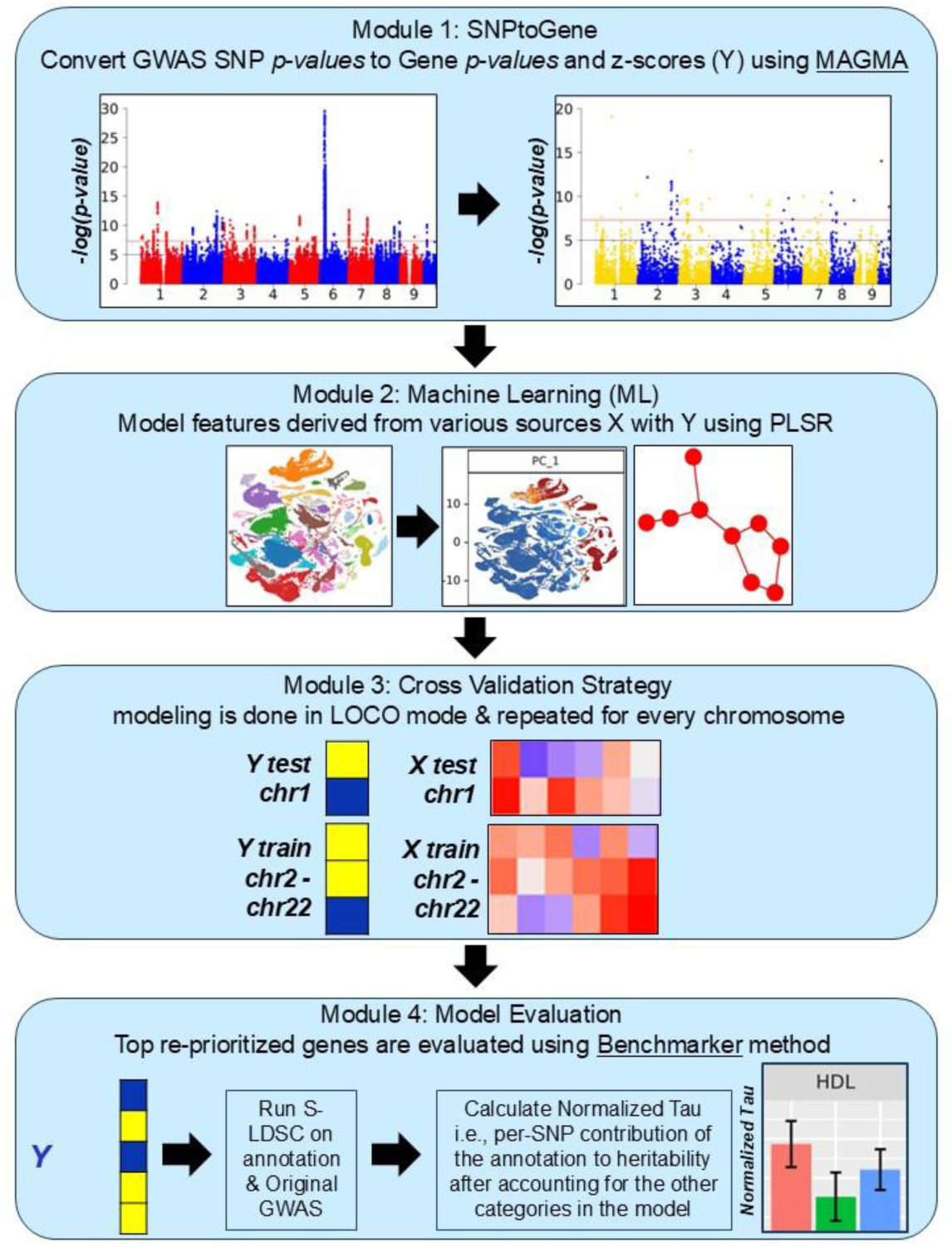
Overview of GMIP Framework. Figure 1: Description GMIP is a comprehensive pipeline designed to integrate GWAS data with multi-omics information. It comprises three key modules: SNPtoGene Module

- **Purpose**: Converts SNP-level GWAS p-values to gene-level p-values and z-scores.
- **Method**: Utilizes MAGMA, a tool that applies multiple regression models to aggregate SNP data to the gene level. Machine Learning (ML) Module

- **Purpose**: Models gene-level z-scores using features derived from diverse sources.
- **Method**: Employs Partial Least Square Regression (PLSR) with features from scRNA-seq, Protein-Protein Interactions (PPI), pathways, etc. The Leave One Chromosome Out Cross-validation (LOCO-CV) strategy is used to evaluate the model’s performance. Model Evaluation Module

- **Purpose**: Assesses the performance of the gene prioritization from the ML module. **Method**: Uses the benchmarker strategy, where top N re-prioritized genes are annotated and evaluated with S-LDSC (LD Score Regression) to calculate normalized Tau scores for each top gene list.

### 2.2 GWAS Studies Used for GMIP Evaluation

We utilized 8 different GWAS studies for initial evaluation of GMIP’s parameter performances, selected for their diverse phenotypes and high heritability, sufficient sample sizes, and large number of significant SNPs and loci (Table S1). For expanded analysis, we calculated heritability across 319 public domain traits using LDSC (Bulik-Sullivan et al., 2015) and selected 46 that represented a broad range of heritability values (Table S2 & S3).

### 2.3 Gene Features from Multi-Omics Datasets

In GMIP, we utilized gene features generated and used by the NetWAS (C. S. Greene et al., 2015), NAGA (Carlin et al., 2019; Huang et al., 2018), and PoPS studies (Weeks et al., 2023). All networks were preprocessed to have ENSEMBL (Harrison et al., 2024)gene identifiers to maintain consistency across comparisons. Briefly, the features were derived as follows:

#### NetWAS Networks as Features

The NetWAS study constructed tissue-specific networks by integrating 987 genome-scale datasets, including expression and interaction data from BioGRID (Oughtred et al., 2021), IntAct (Kerrien et al., 2012), MINT (Licata et al., 2012), and MIPS(Mewes et al., 1999). Interactions were categorized based on experimental evidence, and transcription factor binding was assessed using JASPAR (Rauluseviciute et al., 2024) motifs via FIMO (Grant et al., 2011)within 1 kb upstream of genes, represented as binary scores. Gene expression data from GEO was processed into Pearson correlations, and a naïve Bayesian classifier estimated gene interactions across 144 tissues. The final network included four edge classes (C1-C4) for tissue specificity and used the Sleipnir library in C++. A tissue-naïve global network from the NetWAS study is included in GMIP.

#### NAGA PCnet network as Features

The Parsimonious Composite Network (PCNet) was constructed by combining multiple networks, such as protein-protein interactions, gene co-expression, genetic interactions, pathways, and literature-derived networks. Initially, each network’s ability to predict biological functions was assessed, and various integration methods were tested (Huang et al., 2018). The final PCNet required support from at least two networks per interaction, outperforming the best individual network (STRING (Szklarczyk et al., 2023)) while being smaller in size. This network was then used in the NAGA study and integrated into GMIP.

#### PoPS Features

○ **Bulk or scRNA-seq datasets:** 77 gene-expression datasets were standardized using Seurat v.3. Cells were filtered based on quantile distribution, data was then scaled, and principal components (PCs) and gene loadings were computed using advanced statistical methods like PCA, ICA etc. Clusters were identified using the Louvain algorithm, visualized with UMAP, and adjusted for batch effects when needed. Differential expression was carried out between clusters using a one-versus-all approach with a two-sided Welch’s *t*-test. The detailed pipeline is available here (https://github.com/FinucaneLab/gene_features). PoPS features were then derived across the entire dataset, within specific clusters, and between clusters, focusing on gene expressions, gene loadings, and differential expression statistics to identify significant gene regulation.
○ **Curated biological pathways:** Features were encoded as binary indicators for gene membership in pathways from databases like KEGG, GO, Reactome etc., as curated by DEPICT (Pers et al., 2015).
○ **Predicted PPI networks:** For every gene, its first-degree neighbors in predicted InWeb_IM PPI network (Li et al., 2017) were encoded as a binary indicator to generate features.

For all the above features, a control feature for each dataset was included. Each feature was then standardized to have a mean of 0 and a variance of 1 across all genes. In GMIP we have included the expression, pathway and PPI features each and also a fullset which includes all the three features into a single set.

### 2.4 Feature Construction from NAFLD scRNA-seq Data

In this section, we describe how the NAFLD-specific scRNA-seq data was processed to generate a feature set similar to the general features used in PoPS. The primary objective of this process was to generate biologically meaningful features from the raw expression data, rather than directly using simple gene counts. These scRNA-seq features provide a more nuanced representation of gene activity in specific cell types and functional contexts, which can be crucial for understanding complex traits like non-alcoholic fatty liver disease (NAFLD).

#### A. Data Preprocessing

We began by downloading the NAFLD scRNA-seq dataset (GSE166504) and its metadata from the Gene Expression Omnibus (GEO). Raw gene expression counts were normalized and scaled using the Seurat toolkit, ensuring that highly expressed genes did not dominate the downstream analysis. This normalization was followed by variable gene selection, where the top 2,000 highly variable genes were chosen for further analysis.

#### B. Dimensionality Reduction

Dimensionality reduction techniques like PCA (Principal Component Analysis) and ICA (Independent Component Analysis) were employed to compress the high-dimensional gene expression data into a set of principal components, capturing 80% of the variance. This step ensured a more manageable dataset for feature extraction and prevented overfitting in the later stages.

#### C. Clustering and Feature Generation

We used UMAP (Uniform Manifold Approximation and Projection) and the Louvain clustering algorithm to identify clusters within the data, based on the cell expression profiles. Differential expression analysis between clusters was conducted to derive cluster-specific marker genes, which were then used as features. This approach captures cell-type-specific expression patterns relevant to NAFLD. Key feature types include cluster-based aggregated expression values, differential expression between clusters, and global gene expression levels across the entire dataset. Control features were also created for genes with zero counts across certain datasets to ensure they were treated appropriately in subsequent analyses.

This detailed construction of features ensured that biologically meaningful patterns were incorporated into the PoPS or GMIP-PLSR pipeline for the NAFLD GWAS analysis.

### 2.5 Module 1: Obtaining Gene-Level Scores from GWAS Summary Statistics

We used MAGMA (Multi-marker Analysis of GenoMic Annotation) to convert SNP level p-values from GWAS to gene level p-values and z-scores. MAGMA employs a multiple regression model for gene analysis which accounts for LD and improves power to detect multi-marker associations (de Leeuw et al., 2015). We used 1000 Genomes Phase 3 (Auton et al., 2015) data for EUR individuals to estimate LD. A 0-kb window around the gene body was used to map SNPs to genes like previous studies (Fine et al., 2019; Weeks et al., 2023). MAGMA model then projects the SNP matrix for a gene onto its principal components (PCs), using the linear regression model

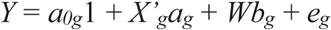

where *X’_g_* is the matrix of PCs, *Y* is the phenotype, *W* is an optional matrix of covariates, *a_g_* represents the genetic effect, *b_g_* the effect of the covariates, *a_0g_* the intercept, and *e_g_* the vector of residuals. In our analysis, the *snp-wise=mean* gene analysis model that uses sum of *-log (SNP p-value)* as test statistic was used. It is more attuned to the mean SNP association, though it skews towards associations in areas of higher LD in a gene. The gene P value is then converted to a z-score using the probit function. This z-score is used in GMIP as the response variable in regression.

### 2.6 Module 2: Machine Learning Methods

#### Network-Assisted Genomic Association (NAGA) Framework

NAGA (Carlin et al., 2019) employs network propagation (Cowen et al., 2017) to diffuse gene scores across the molecular network, implicating nearby genes through association. Using the random walk with restart model (Vanunu et al., 2010), gene scores *F*(*t*) are updated iteratively according to the equation

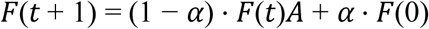

where *⍺* is the propagation constant set by a linear model estimated by network density (Huang et al., 2018), *F*(0) represents the initial gene mutation profile, and *A* is the degree-normalized adjacency matrix. As *t* → ∞, the model converges, producing a reprioritized gene list that incorporates both gene-specific mutations and network-wide associations.

#### Polygenic Priority Score (PoPS) Method

PoPS (Weeks et al., 2023)operates under the assumption that causal genes share functional characteristics. The method begins by using MAGMA to calculate gene-level association statistics from GWAS summary data, considering linkage disequilibrium (LD) and, optionally, covariates like gene length (in this analysis, covariates were not included).

##### Step 1: Feature Selection

Marginal feature selection is performed like MAGMA, where a gene feature *X*_*f*_ is tested for enrichment using a generalized least squares (GLS) model:

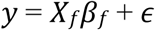

where *y* is the gene-level association, *β*_*f*_is the effect size of the feature, and *ε* represents error terms assumed to follow a multivariate normal distribution (MVN) with a covariance matrix *R* that accounts for LD between genes. Features passing the significance threshold (*P*<0.05) are retained.

##### Step 2: Joint Model Fitting

After feature selection, PoPS fits a joint model by replacing *X*_*f*_ with a matrix *X*, which contains all selected features:

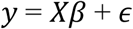

PoPS extends the GLS model by incorporating L2 regularization (ridge penalty) to handle multiple features and improve generalization in the test set.

##### Step 3: Polygenic Priority Score (PoP Score)

PoPS computes a polygenic priority score (PoP score) for each gene *g* using:

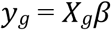

Here, *X*_*g*_ represents the feature set for gene *g*, and *β* is the estimated effect sizes from the joint model. A gene is prioritized if it is in a 1-Mb locus centered on a genome-wide significant variant and has the highest PoPS score in that region.

#### Regressing GWAS Gene Scores on Gene Features using PLSR (GMIP-PLSR)

The GMIP-PLSR pipeline is designed to enhance PoPS by incorporating Partial Least Squares Regression (PLSR) for more effective modeling. After PoPS processes the training data, GMIP-PLSR utilizes PLSR to address the issue of multicollinearity and improve the predictive accuracy of the models. PLSR is particularly well-suited for this task as it simultaneously reduces dimensionality while maximizing the covariance between predictors (gene features) and responses (GWAS z-scores).

PLSR is a dimensionality reduction technique that builds predictive models by extracting latent variables (LVs) from both the predictors (gene features) and response (GWAS z-scores) matrices (Ortiz et al., 2023). These LVs are linear combinations of the original variables that capture the maximum covariance between the two datasets. PLSR effectively handles collinear predictors, a common challenge in omics studies where gene features are often highly correlated. While ridge regression can also handle collinearity by penalizing large regression coefficients, PLSR offers several advantages:

##### A. Dimensionality Reduction

PLSR simultaneously reduces the dimensionality of both predictors and responses, leading to a more parsimonious model compared to ridge regression, which only addresses the predictor space (Carrascal et al., 2009; Wold, 1975).

##### B. Latent Variable Interpretation

The extracted LVs often have biological interpretations, providing insights into the underlying relationships between genetic variants and the phenotype. Ridge regression does not offer such interpretability (Lê Cao et al., 2009; Nguyen & Rocke, 2002).

##### C. Predictive Performance

In many cases, PLSR has been shown to outperform ridge regression in terms of predictive accuracy, especially when dealing with highly correlated predictors, as is common in GWAS studies (Boulesteix & Strimmer, 2007; Frank & Friedman, 1993).

PLSR seeks to find latent variables (T) and (U) that maximize the covariance between the predictor matrix X (gene features) and the response matrix Y (GWAS z-scores) subject to the constraint that the LVs have unit variance. The model can be expressed as:

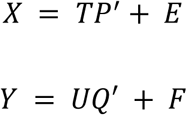

Where *X* is the *n x p* predictor matrix, *Y* is the *n x q* response matrix, *T* and *U* are the latent variable scores *(n x k)*, *P* and *Q* are the loading matrices (*p x k* and *q x k*, respectively), *E* and *F* are the residual matrices. PLSR iteratively identifies the optimal LVs using the Nonlinear Iterative Partial Least Squares (NIPALS) algorithm. Selecting the correct number of components (nc) is crucial, as too few LVs can underfit the model, while too many can lead to overfitting. For this study, we tested two and three components across smaller subset of GWAS datasets and expanded to a range of 2-10 components for larger set. Optimal results were typically observed with three components (nc=3), in line with findings from other studies (Mbatchou et al., 2021; Ortiz et al., 2023).

### 2.7 Module 3: Cross-Validation Strategies

#### A. noCV: Not using any Cross Validation

In GMIP it is possible to completely turn off cross validation and mimic the execution of original NetWAS, NAGA & PoPS studies. In this case the full set of features and gene scores are used for training and testing. Usually, such models are highly biased and not generalizable (Dietterich, 1998). It has been shown that even though results from such models may perform well in gold standard evaluations but the probability of discovering novel relationships is very low (Fine et al., 2019).

#### B. Stratified k-Fold Cross Validation strategy

None of the studies evaluated by us have used k-Fold Cross Validation, but we still include it in GMIP as it is a very popular and effective CV strategy widely used in various ML tasks. Since our GWAS gene scores are highly imbalanced in the sense that there are much fewer significant associations we decided to perform stratified k-fold CV. We first binarized the GWAS gene scores using a common and effective cut off 5e-8 and then performed stratified k-fold cross validation. This is a flexible parameter in the pipeline and can be varied as needed.

#### C. Leave One Chromosome Out Cross Validation strategy (LOCO-CV)

For all our analysis we adapted the Leave One Chromosome Out (LOCO) Cross Validation strategy. In this strategy we leave a chromosome out as testing chromosome and perform all the projections, feature selections and regressions on the other chromosomes. Models made from training chromosomes are then applied to the left-out testing chromosome and scores predicted for each gene on this test chromosome are recorded. By iteratively leaving out each of the 22 autosomes in the human genome, a complete list of predicted test gene scores can be obtained. This list is then sorted by the largest predicted score, and top N genes are chosen for further analysis. Using LOCO ensures that any information leakage due to chromosomal proximity between genes and gene features does not occur between training and testing sets (Dietterich, 1998; Mbatchou et al., 2021; Qian et al., 2023; Rabinowicz & Rosset, 2022).

### 2.8 Module 4: Model Evaluation Strategies

#### A. Using benchmarker strategy for evaluation

Benchmarker (Fine et al., 2019)is an unbiased, data-driven strategy to evaluate gene prioritization algorithms. The method assumes that Single Nucleotide Polymorphisms (SNPs) located near causal genes (genes influencing the trait) are more likely to contribute to the heritability of the trait. Benchmarker uses Stratified Linkage Disequilibrium Score Regression (S-LDSC) to estimate the mean influence of SNPs adjacent to the re-prioritized genes on per-SNP heritability. This analysis considers the specific genes of interest in conjunction with 53 other annotations included in a “baseline model.” This baseline model encompasses various functional genomic regions, such as genic regions, regulatory elements, and conserved sequences. The performance of the chosen genes is assessed using the regression coefficient (*tau*) and its associated p-value under the assumption *tau* > 0. Here, *tau* signifies the impact of SNPs near the prioritized genes on per-SNP heritability, factoring in the baseline annotations. For consistency across different traits, *tau* is normalized by the average per-SNP heritability for each trait, a measure referred to as normalized *tau*. Comparing GWAS re-prioritization results using original GWAS without any gold standard is an important strategy highlighted by the Benchmarker method. GMIP can easily produce prioritized gene lists using LOCO-CV strategy and can be readily run with the benchmarker method. In our analysis we used the top 500 genes after compiling the entire re-prioritized list as in PoPS article. We controlled multiple hypothesis testing and applied the Bonferroni correction to the enrichment p-values.

#### B. Gene Set Enrichment Analysis (GSEA) for Evaluating Gene Reprioritization

Gene Set Enrichment Analysis (GSEA) (Subramanian et al., 2005)is a statistical method used to determine whether a predefined set of genes (gene set) is overrepresented at the top (or bottom) of a ranked list of genes compared to what would be expected by chance. In GMIP, we use the top 500 genes identified by MAGMA from the original GWAS as the predefined gene set *S*. Simultaneously, genes are ranked based on predicted scores from methods like PoPS or NAGA, resulting in a ranked list *L*={*g*1,*g*2,…,*gN*}, where *g* represents genes sorted by their predicted scores. GSEA computes an Enrichment Score (ES), which reflects the concentration of the gene set *S* within this ranked list by calculating the deviation of the observed rankings from what would be expected by random chance. The ES measures the extent to which the genes in the set *S* are enriched at the top of the ranked list. To determine the significance of the ES, GSEA compares it to a distribution of enrichment scores generated from random permutations of the gene labels, producing a Normalized Enrichment Score (NES). The significance of the NES is then assessed using a Kolmogorov-Smirnov (KS) test to compute a p-value. Since the purpose of GMIP is to enhance GWAS results, we expect that significant genes identified in the original GWAS should still be found within the top portion of the reprioritized list. By using GSEA, we evaluate whether the original significant GWAS genes are enriched at the top of the reprioritized gene list, confirming the effectiveness of the reprioritization performed by GMIP.

### 2.9 Multicollinearity Analysis and PoPS Feature Clustering

While PoPS marginal OLS effectively selects features based on their association with GWAS gene z-scores, multicollinearity among the selected features remained an issue. This was initially hinted at by PoPS’ post-reprioritization clustering analysis, which demonstrated some degree of redundancy among features. To confirm this, we assessed multicollinearity within the predictor matrix by calculating the Condition Index (CI) for each selected predictor, which quantifies the severity of multicollinearity. The CI is calculated by first constructing the cross-product matrix (*X*′*X*) of the predictor matrix. This matrix represents the interactions between the predictor variables. We then perform eigenvalue decomposition on the *X*′*X* matrix, as the eigenvalues provide insight into the linear dependencies within the predictor set. To avoid issues from near-zero or negative eigenvalues due to numerical instability, we filter out eigenvalues below a small threshold (1e-10). The CI is then calculated as the square root of the ratio between the largest and each individual eigenvalue, where high CI values (>30) indicate multicollinearity (Belsley et al., 1980; W. H. Greene, 2003). Multicollinearity reduces the interpretability of regression models and can inflate the standard errors of coefficients, leading to unreliable results. Despite Ridge regression’s inherent ability to handle correlated features to a certain extent, the magnitude of multicollinearity we observed (Figure 2) indicated that more sophisticated techniques were needed to optimize the feature set for modeling.

**Figure 2:**
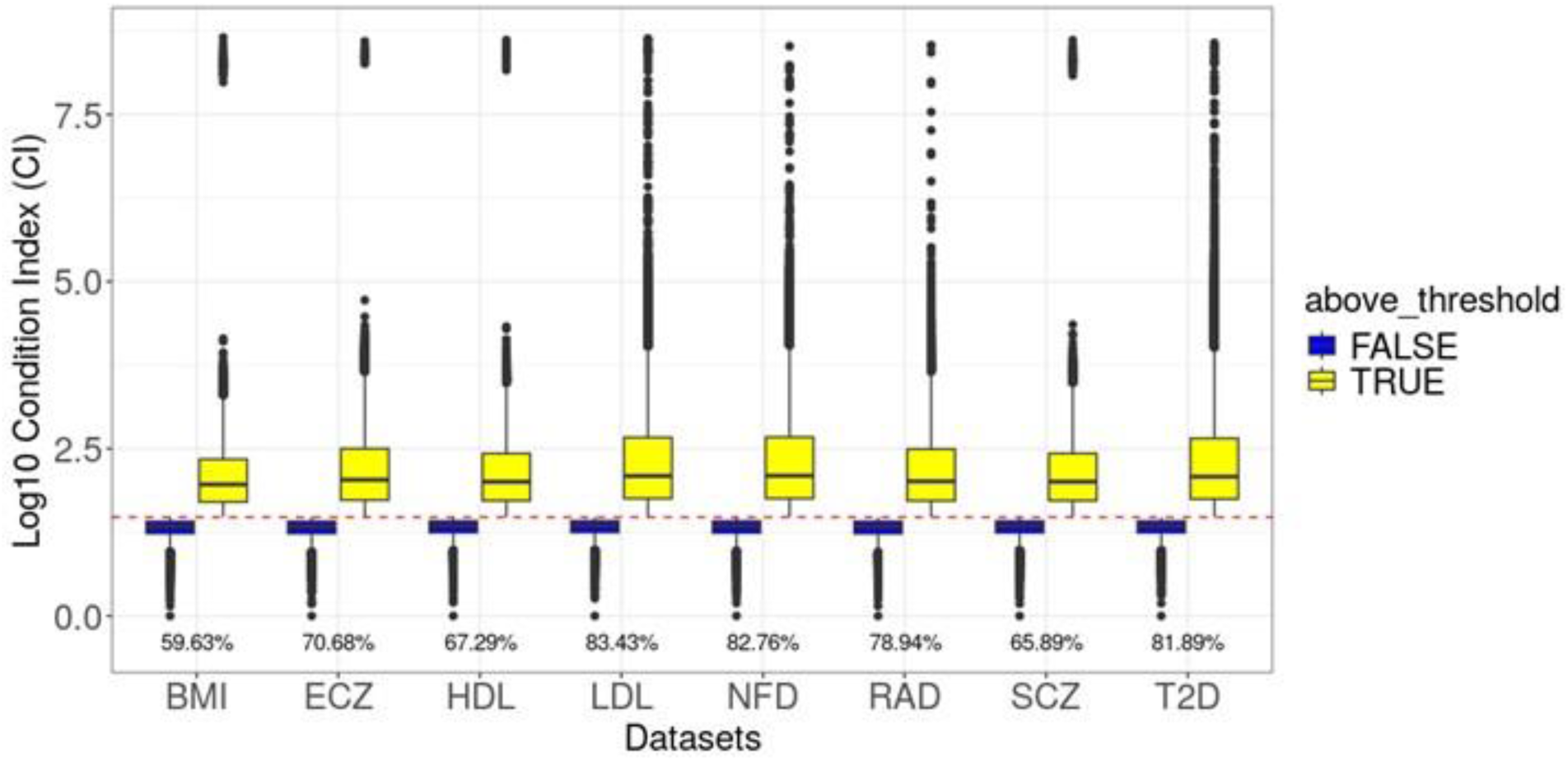
Condition Index (CI) distribution for features selected by marginal OLS. Figure 2: Description The box plot shows the distribution of log-transformed Condition Index (CI) values across eight datasets (BMI, ECZ, HDL, LDL, NFD, RAD, SCZ, and T2D).

- The dashed red line indicates the CI threshold of 30, where multicollinearity becomes problematic.
- Yellow points represent features with CI values greater than 30, while blue points represent values below.
- Percentages below each dataset reflect the proportion of features with CI values exceeding 30. High multicollinearity persists in all datasets potentially impacting PoPS ridge regression.

### 2.10 Techniques for Addressing Multicollinearity

We explored three primary approaches to mitigate multicollinearity:

#### Feature Clustering with PCA and Ridge Regression

This method builds on the post-reprioritization strategy used in PoPS, with additional refinement steps to enhance feature selection. We first applied Incremental Principal Component Analysis (IPCA) to the feature matrix, allowing us to handle large datasets in a memory-efficient manner while capturing the majority of the variance—typically set to 80%. Once PCA reduced the dimensionality, we used hierarchical clustering on the principal components to identify groups of correlated features. Using complete linkage as our clustering method, we evaluated the quality of the clustering with the silhouette score. For each cluster, the most predictive feature was selected based on the marginal OLS beta coefficient from PoPS. The feature with the highest absolute beta value in each cluster was retained for modeling, which reduced the number of redundant features by approximately 30-40% across the initial eight GWAS datasets. The refined feature set was then passed to Ridge regression, like the approach in PoPS, but with reduced collinearity among the features.

#### Dimensionality Reduction via PCA Alone

In this approach, we sought to reduce the dimensionality of the feature set without performing any clustering, relying solely on Principal Component Analysis (PCA) to simplify the feature space. PCA alone was applied to the complete set of selected features to explain 80% of the variance in the data. By transforming the features into uncorrelated principal components, this method directly addresses the issue of multicollinearity without retaining the original feature structure. The principal components generated are linear combinations of the original features, and each component represents a portion of the total variance in the dataset. Once the dimensionality was reduced, we used the transformed principal components as inputs for Ridge regression.

#### Partial Least Squares Regression (PLSR) with Different Numbers of Components

Given the encouraging results from dimensionality reduction techniques, we explored PLSR, a method that directly addresses multicollinearity by constructing latent variables that capture the covariance between predictors and response matrices. Initially, we tested PLSR with two and three components, finding that PLSR with three components (nc3) performed best across most GWAS datasets. For the RAD trait, normalized tau values improved dramatically from 2.9984 (PoPS) to 5.0183 using PLSR. In BMI, the normalized tau score increased from 0.2618 to 0.3893, and for LDL, it rose from 5.0759 to 5.2691. PLSR consistently outperformed both PoPS and PCA + ridge regression, highlighting its robustness in addressing multicollinearity and improving GWAS gene prioritization (Figure 3).

**Figure 3:**
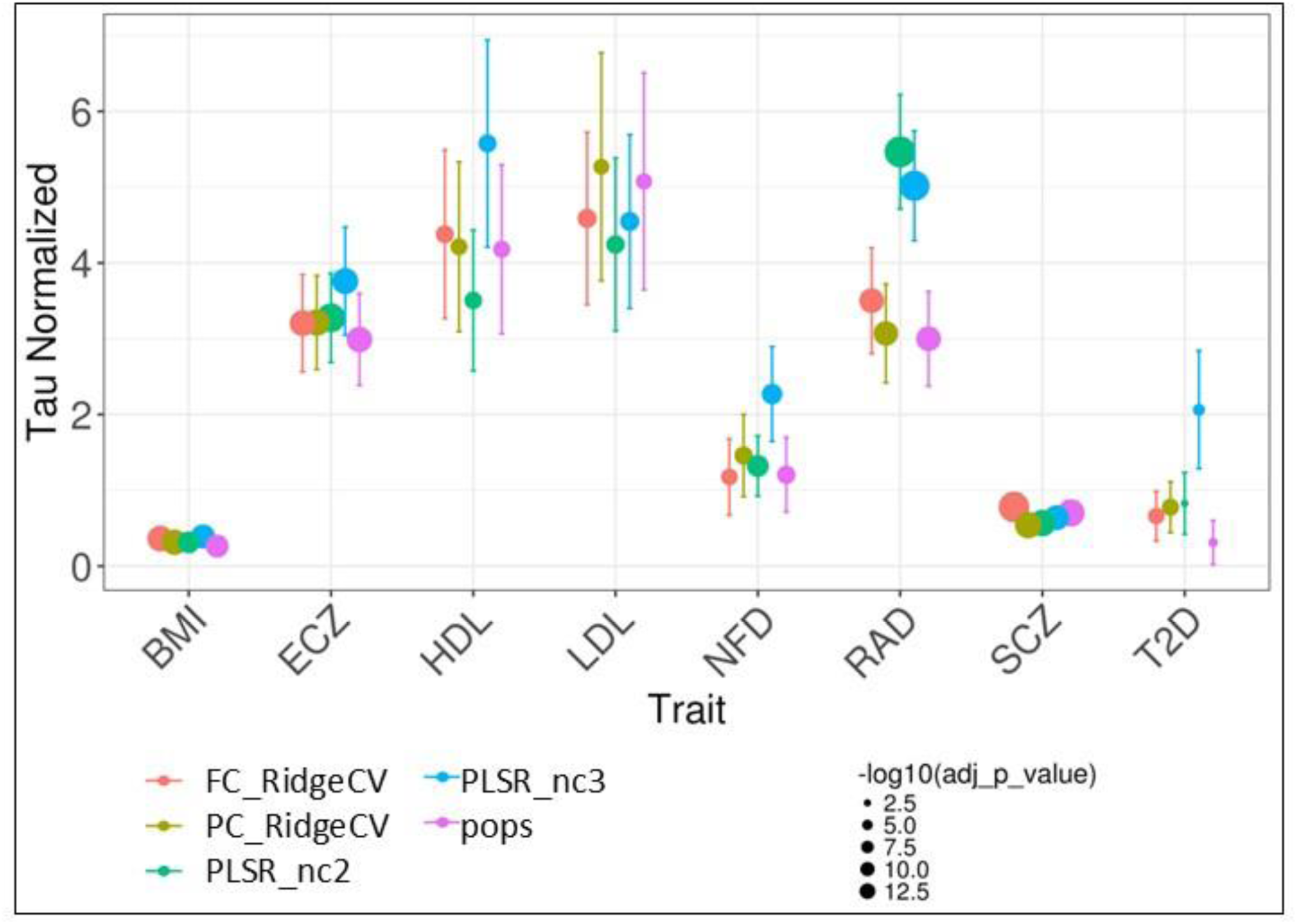
Tau Score Comparison Across Different Methods for various GWAS. Y-axis represents Normalized tau values representing the per-SNP heritability explained by the top 500 re-prioritized genes. X-axis represents eight GWAS traits. The size of points indicates the statistical significance of the enrichment (larger points = more significant results). Error bars represent the standard error of the tau scores for each method. Methods compared:

- FC_RidgeCV (Feature Clustering with RidgeCV)
- PC_RidgeCV (Dimensionality Reduction via PCA with RidgeCV)
- pops (PoPS Ridge Regression)
- PLSR_nc2 (Partial Least Squares Regression with 2 components)
- PLSR_nc3 (Partial Least Squares Regression with 3 components) **Key results:** Methods with dimensionality reduction score better than ridge regression in PoPS. PLSR_nc3 performs better for most GWAS compared to other methods.

## 3.0 Results

### 3.1 Overview of GMIP Framework and Optimal Module Selection

GMIP is structured around four essential modules (Figure 1): SNP2Gene Mapping, Machine Learning Modeling, Cross-Validation Strategy, and Evaluation. This modular pipeline provides flexibility, allowing users to explore various combinations by incorporating multiple choices for each module. We initially tested each method using the original study’s module settings. Afterward, we kept the machine learning module constant and varied other module choices to evaluate performance changes. This approach was applied to both the NAGA and PoPS studies.

In Module 1 (SNP2Gene Mapping), MAGMA emerges as the preferred tool for handling linkage disequilibrium (LD) and performing robust gene-level association analysis. Module 2 (Machine Learning) leverages PoPS with ridge regression, which demonstrates reliable performance across various GWAS datasets. For features, both NAGA and PoPS feature sets perform well, but PoPS features are preferred due to their superior interpretability in associating gene pathways, gene-cell types, and GWAS traits. In Module 3 (Cross-Validation Strategy), the Leave-One-Chromosome-Out (LOCO-CV) method is used to prevent information leakage, a common issue in k-fold cross-validation due to chromosomal proximity. Finally, in Module 4 (Evaluation), Benchmarker is utilized as the most sensitive strategy, providing key insights into the heritability of reprioritized genes. These optimal module choices are illustrated within the overall framework in Figure 1, forming the foundation for applying GMIP across different GWAS studies, allowing us to address multicollinearity and explore advanced regression techniques, including Partial Least Squares Regression (PLSR).

Comprehensive benchmarking indicated that the combination of PoPS features, Ridge regression, LOCO-CV, and Tau heritability evaluation performed consistently well across traits (Supplementary Figure S2). Based on these findings, we kept the framework fixed and replaced Ridge with PLSR to mitigate multicollinearity and further improve gene prioritization. We began by evaluating the performance of the original GMIP pipeline using PoPS features and Ridge regression, which revealed substantial multicollinearity. This motivated us to test dimensionality reduction and regression alternatives, including PCA and PLSR, as detailed below.

### 3.2 Evaluation of Gene Reprioritization Tools within GMIP

For all GWAS studies, NAGA performed well when using either of the gene-gene feature sets (PCNet & NetWAS) without cross-validation (noCV) (Figure S1). This was evident in the GSEA of top 500 genes (based on training MAGMA z-scores) in gene list ranked by NAGA predicted scores. These noCV results are consistent with what is reported by the NAGA study. However, performance dropped significantly with LOCO-CV or kFOLD-CV, indicating potential overfitting. Normalized Tau scores calculated for top 500 genes predicted by NAGA models using LOCO-CV were also low for most GWAS, except for RAD. This suggests that normalized tau and GSEA scores are correlated and provide complementary insights.

Like NAGA, PoPS performed well without cross-validation (noCV) across all features (Figure S2), as shown by the GSEA analysis. However, when LOCO-CV or kFOLD-CV was applied, PoPS maintained significant GSEA results, indicating it is better at building generalizable models than NAGA. Normalized Tau scores calculated by Benchmarker for the top 500 genes predicted by PoPS, using a combination of NAGA features (PCNet) and full PoPS features, were also high and significant for most GWAS datasets.

### 3.3 Multicollinearity Between Selected Features

After projecting out gene-level covariates using MAGMA, features selected via marginal OLS in PoPS were passed into the GMIP pipeline for joint modeling. Significant multicollinearity was observed across all eight GWAS datasets, as indicated by the high percentage of selected predictors with a Condition Index (CI) greater than 30 (Figure 2). These predictors, derived from a diverse set of omics features—including scRNA-seq datasets, pathway memberships, and protein-protein interaction networks—exhibited strong inter-feature correlations. This multicollinearity impaired ridge regression’s capacity to effectively model the relationships between predictors and GWAS z-scores, leading to inflated standard errors of regression coefficients and reduced interpretability. Although ridge regression can handle some degree of multicollinearity, the extent observed in this analysis significantly degraded its performance. To address this, we explored dimensionality reduction methods like Principal Component Analysis (PCA) and Partial Least Squares Regression (PLSR), which are well-suited for managing multicollinearity by creating latent variables that encapsulate the maximal variance within the predictor space. These techniques improved the model’s ability to handle collinear predictors, enhancing interpretability and reducing error variance in the regression coefficients.

### 3.4 Dimensionality Reduction Techniques for Addressing Multicollinearity

To address the multicollinearity observed between the features selected by PoPS, we applied two key dimensionality reduction techniques: Feature Clustering with PCA and Dimensionality Reduction via PCA Alone. Both methods aim to simplify the feature space while preserving predictive performance and improving the downstream regression models.

#### Feature Clustering with PCA and Ridge Regression

This approach combines PCA with hierarchical clustering to group features based on their correlation structure. By reducing redundancy and retaining the most predictive feature from each cluster, we effectively addressed multicollinearity. When applied to the eight initial GWAS studies, we observed a substantial improvement in normalized tau scores. For instance, in the BMI dataset, the normalized tau score increased from 0.2618 (PoPS) to 0.3893 with feature clustering and PCA. Similarly, for ECZ, the normalized tau score improved from 2.9888 to 3.7598. Other traits, such as HDL and LDL, also showed notable improvements, confirming that addressing multicollinearity significantly enhances model performance (Figure 3).

#### Dimensionality Reduction via PCA Alone

In this method, we reduced the dimensionality of the full feature set using PCA, aiming to explain 80% of the variance without performing feature clustering. The principal components were then used in ridge regression. While some traits, such as BMI (normalized tau score improved from 0.2618 to 0.3137) and HDL (normalized tau score increased from 4.1795 to 4.2133), benefited from this approach, others, like SCZ, showed a decline in performance (normalized tau score decreased from 0.7004 to 0.5405). This suggests that while PCA alone can be helpful in some cases, it may not handle the complex correlations in omics data as effectively as feature clustering (Figure 3).

#### Partial Least Squares Regression (PLSR) with Different Numbers of Components

Given the encouraging results from dimensionality reduction techniques, we explored PLSR, a method that directly addresses multicollinearity by constructing latent variables that capture the covariance between predictors and response matrices. Initially, we tested PLSR with two and three components, finding that PLSR with three components (nc3) performed best across most GWAS datasets. For the RAD trait, normalized tau values improved dramatically from 2.9984 (PoPS) to 5.0183 using PLSR. In BMI, the normalized tau score increased from 0.2618 to 0.3893, and for LDL, it rose from 5.0759 to 5.2691. PLSR consistently outperformed both PoPS and PCA + ridge regression, highlighting its robustness in addressing multicollinearity and improving GWAS gene prioritization (Figure 3).

### 3.5 GMIP Outperforms PoPS for Most GWAS Re-Prioritizations

After establishing the effectiveness of PLSR within GMIP across eight GWAS datasets, we expanded our analysis to a larger set of GWAS datasets, leveraging heritability estimates calculated using LDSC (Bulik-Sullivan et al., 2015). Heritability serves as an important indicator of whether a GWAS summary dataset is suitable for reprioritization. For this expanded analysis, we calculated heritability across 319 public domain traits and selected 46 that represented a broad range of heritability values (Figure 4 & Table S3).

**Figure 4:**
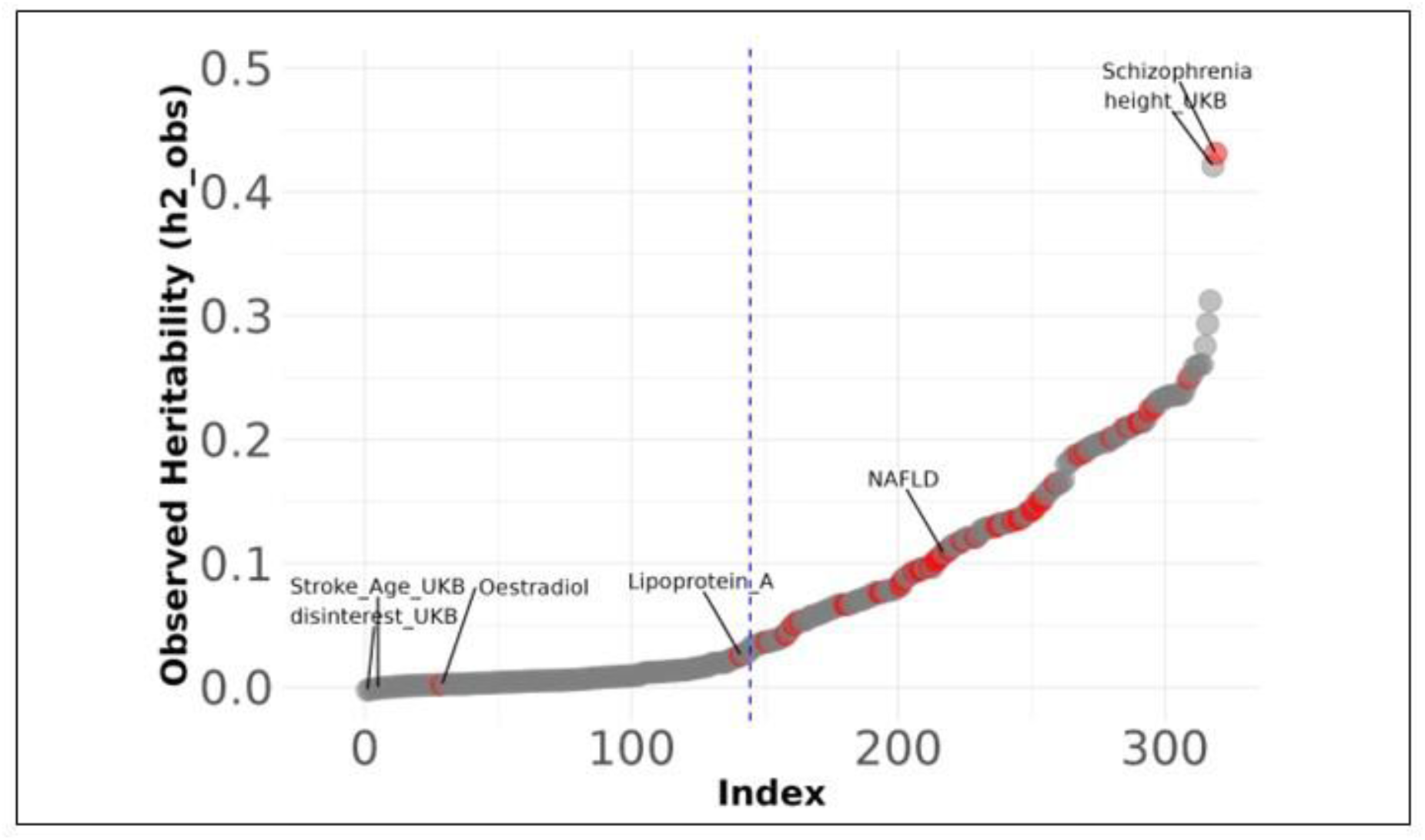
Heritability of Different GWAS Traits. Heritability estimates were derived using LD Score Regression (LDSC) for 319 GWAS traits accumulated from various sources (Table 3.3). A random set of 46 different GWAS (red dots) with a range of observed heritability scores was chosen for deeper analysis (Table 3.2 & 3.3). GWAS traits with observed heritability scores greater than a specific threshold (0.05) were generally successfully re-prioritized using GMIP, leading to significant Tau scores.

Extensive testing was carried out to assess the feasibility of reprioritization for each trait and to determine the optimal hyperparameters and methods. We compared PoPS, which uses ridge regression to model GWAS gene scores, against GMIP-PLSR, which employs Partial Least Squares Regression (PLSR). To evaluate the effectiveness of these approaches, we applied the benchmarker strategy, varying the number of top genes prioritized from GWAS and the number of components used in PLSR.

Our results revealed that 43 out of the 46 GWAS traits could be reprioritized successfully (Table 1), yielding significant tau values (adjusted Enrichment p-value <=0.01). This suggests that GMIP can reliably re-prioritize genes for most traits, even those with varying heritability, and confirms that heritability is a critical factor in determining the utility of reprioritization. Based on these findings, we propose a heritability cut-off for future studies, which can be used to select suitable GWAS datasets for reprioritization within GMIP (Figure 4).

**Table 1:**
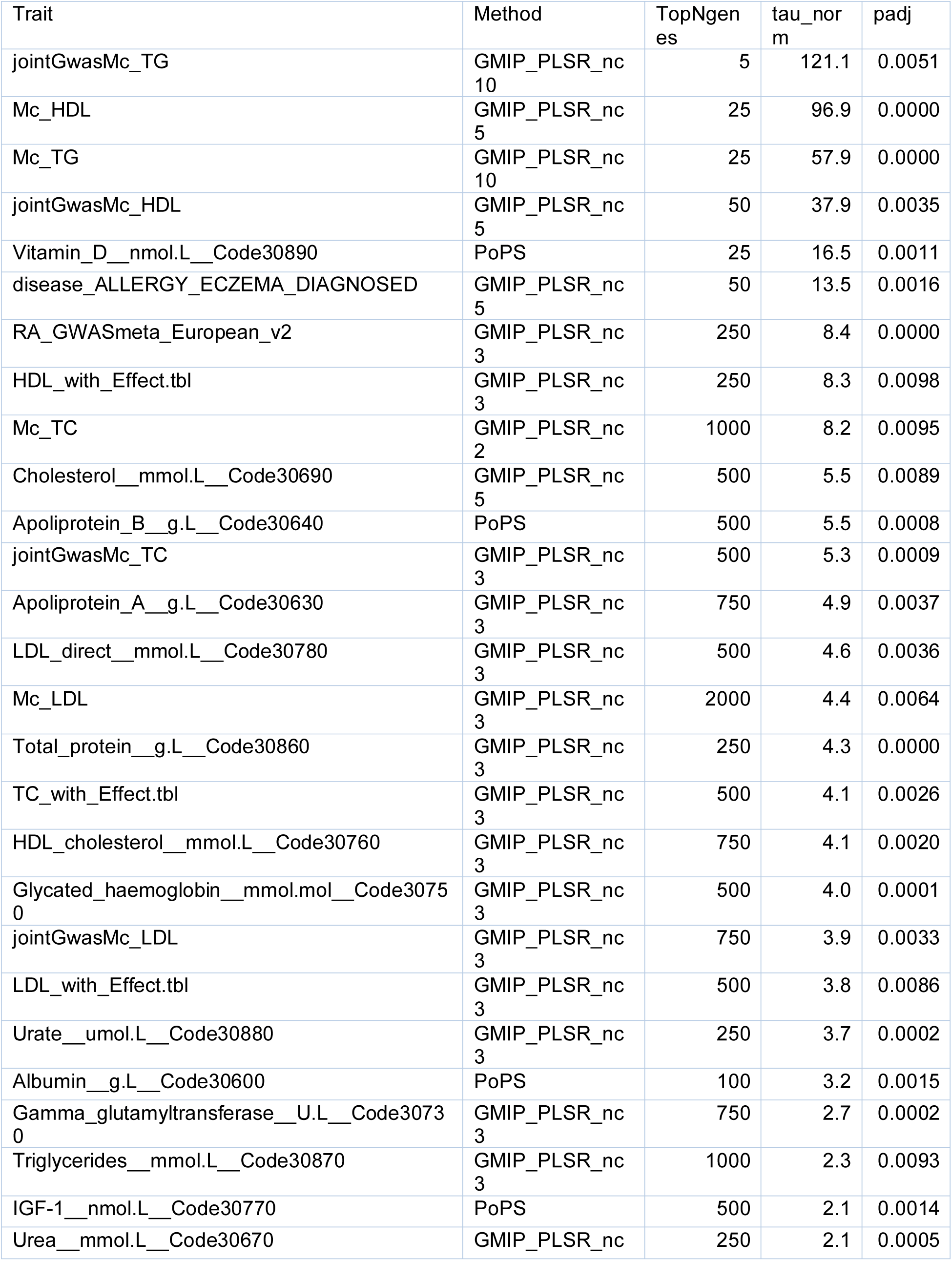
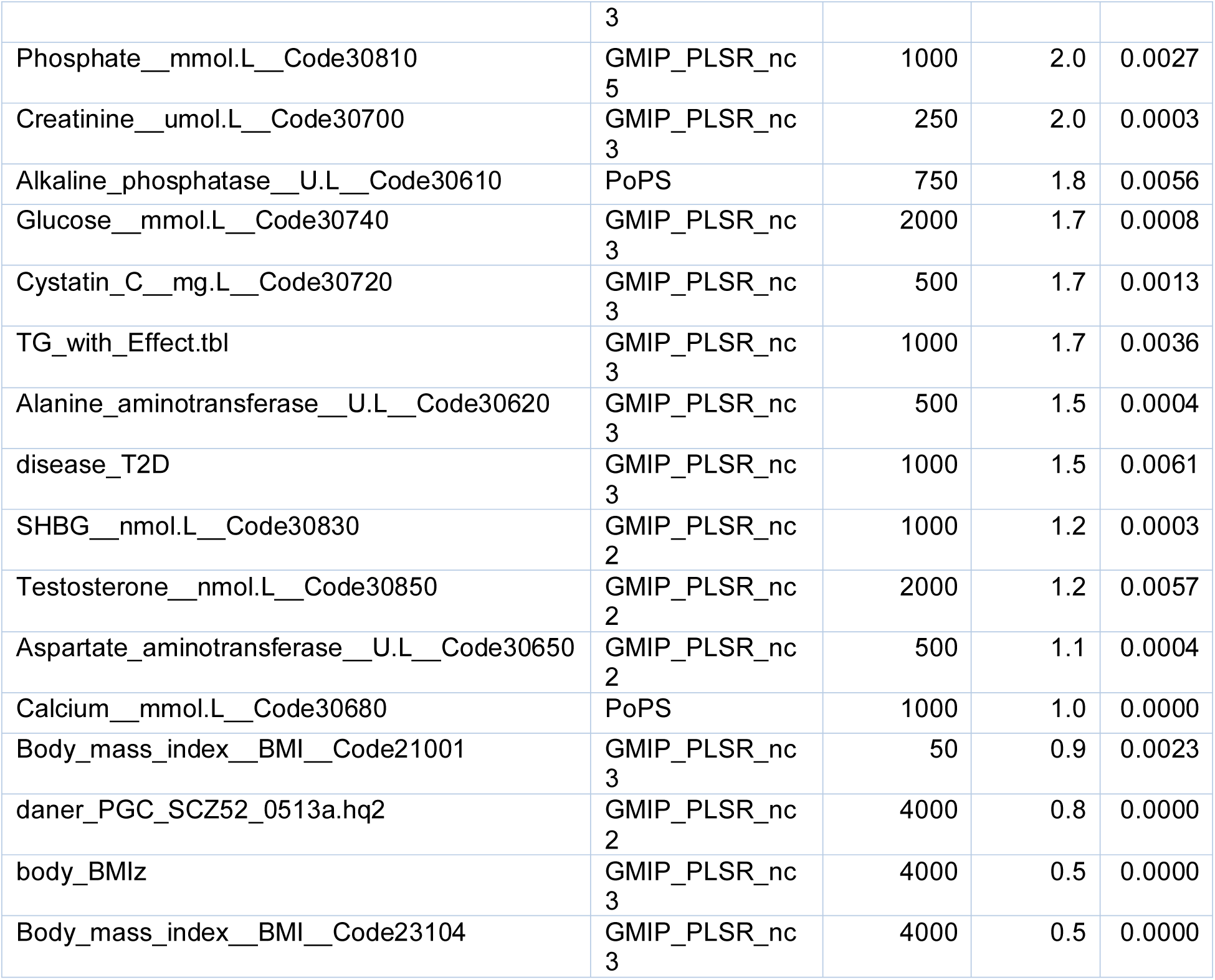
Results of GMIP evaluation for various traits.

In terms of hyperparameter tuning, we found that using the top 500 GWAS genes consistently provided the most significant enrichment and highest tau values. Additionally, varying the number of components in PLSR from 1 to 10 revealed that three components (nc3) performed the best in most cases (Figures 5A & 5B). In nearly all comparisons, GMIP-PLSR with nc3 outperformed PoPS using ridge regression (Figures 5A-B & 6), reaffirming the strength of GMIP-PLSR as a superior method for gene prioritization in diverse GWAS datasets.

**Figure 5:**
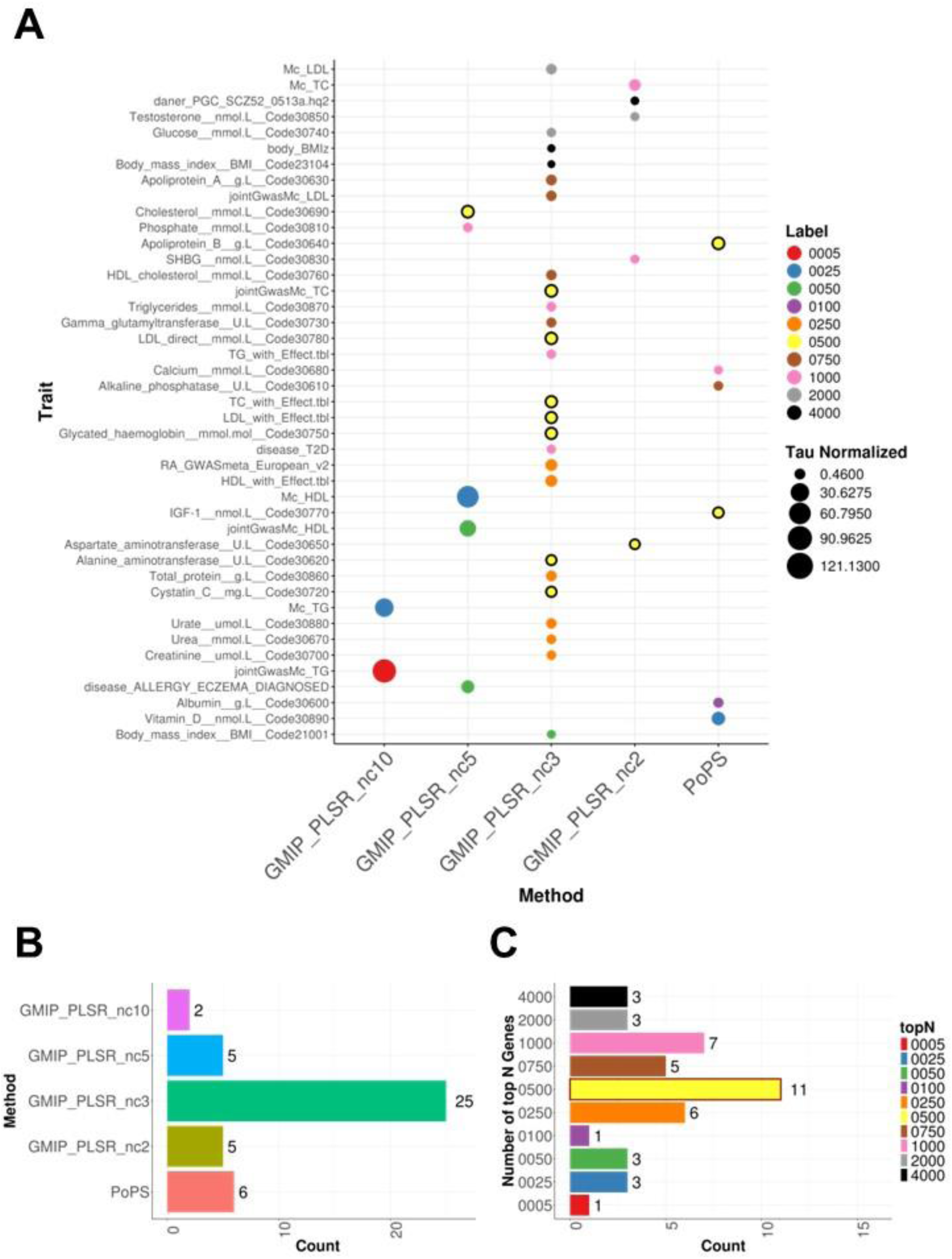
Comparison of GMIP and PoPS with Hyperparameter Optimization. Figure 5: Description **(A)** This panel illustrates the performance of GMIP-PLSR (with varying numbers of components from 2 to 10) and PoPS for 46 different traits (shown on the Y-axis). Both methods were executed in a Leave One Chromosome Out Cross-validation (LOCO-CV) manner. The re-prioritized gene lists, ranging from 5 to 4000 genes, were evaluated using normalized Tau scores calculated through the benchmarker strategy. For each trait (row), the method achieving the highest Tau score and passing the significance threshold of 0.01 for adjusted enrichment p-value (Bonferroni correction) was selected. **(B)** This bar plot shows that GMIP-PLSR with 3 components yielded the best results for the majority of traits, outperforming other configurations and PoPS. **(C)** This bar plot indicates that the top 500 genes consistently provided the most significant and optimal Tau scores across different traits. In summary, GMIP-PLSR demonstrated superior performance in re-prioritizing GWAS traits compared to PoPS, particularly when using 3 PLSR components and focusing on the top 500 genes, as evidenced by the highest normalized Tau scores and significance levels.

**Figure 6:**
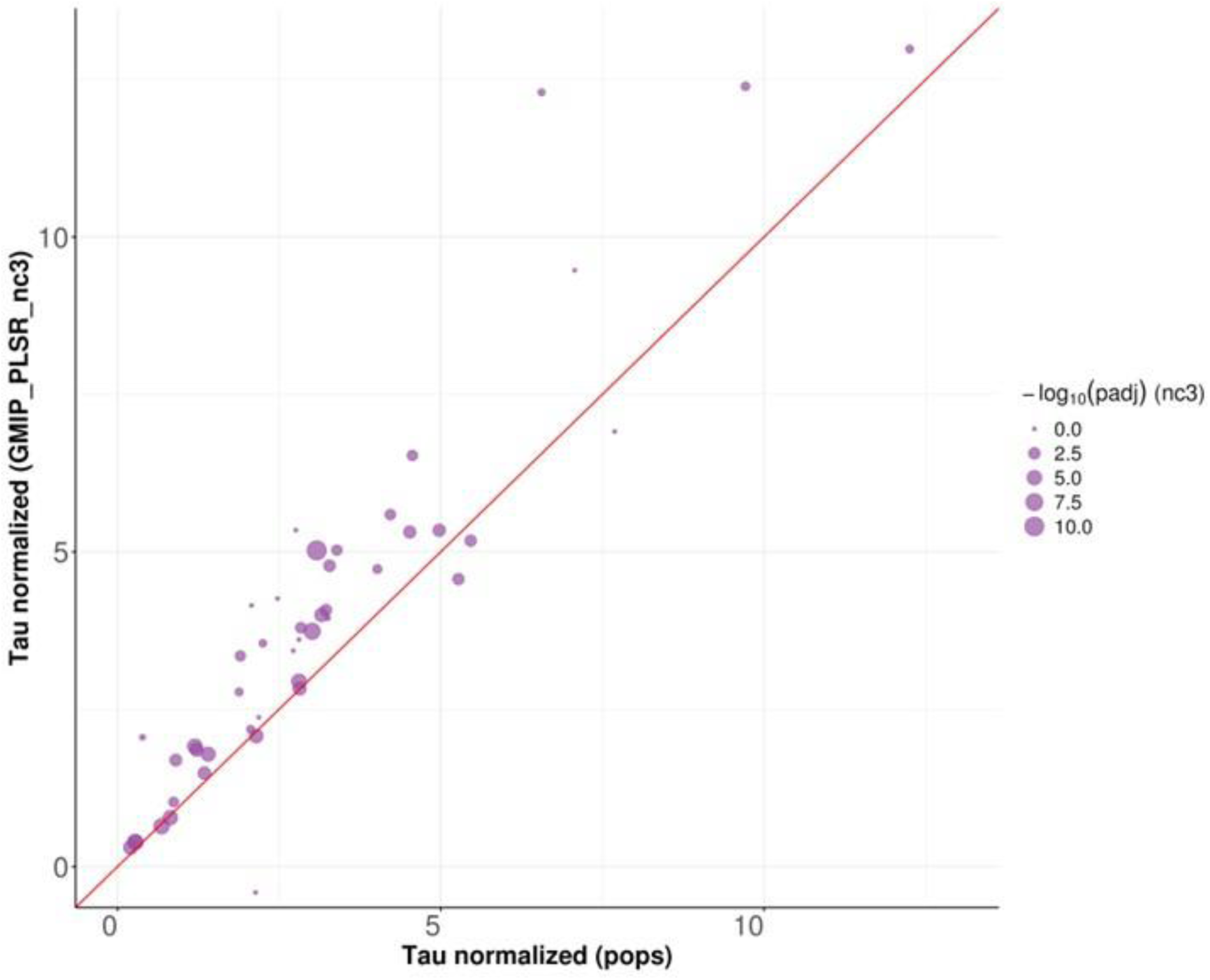
Comparison of Normalized Tau Scores Between PoPS and GMIP-PLSR. This scatter plot compares the normalized Tau scores calculated for two methods: PoPS (X-axis) and GMIP-PLSR with 3 components (Y-axis). Each point represents a different trait, with the size of the points corresponding to the adjusted enrichment p-value for GMIP Tau scores, as calculated by the benchmarker. The red diagonal line represents the line of equality, where points along this line indicate equal performance of both methods. Points above the line suggest better performance of GMIP-PLSR, while points below the line suggest better performance of PoPS. The overall trend, with most points lying above the red line, indicates that GMIP-PLSR generally performs better than PoPS in re-prioritizing GWAS traits.

### 3.6 Comparison of NAFLD GWAS Re-Prioritization Using Various Features

To assess whether disease-specific feature sets outperform general features from public databases like PoPS, we applied the GMIP pipeline to the non-alcoholic fatty liver disease (NAFLD) GWAS. Two feature sets were evaluated: general PoPS features and features derived from a NAFLD-specific mouse single-cell RNA-seq dataset. Our results indicate that the GMIP_PLSR_nc3 model using PoPS features produced the highest tau value (2.96) for the top 250 genes, suggesting the strongest enrichment. In contrast, the same GMIP_PLSR_nc3 model using scRNA-seq-derived features generated a lower yet still significant tau value (1.59) for the top 250 genes.

For consistency, we also included a third set of top 250 genes, ranked directly by MAGMA z-scores from the GWAS analysis. When comparing the overlap between these three sets, significant intersections were observed across all groups (Figure 7), highlighting that each feature set captures relevant biological insights, albeit with varying degrees of specificity.

**Figure 7:**
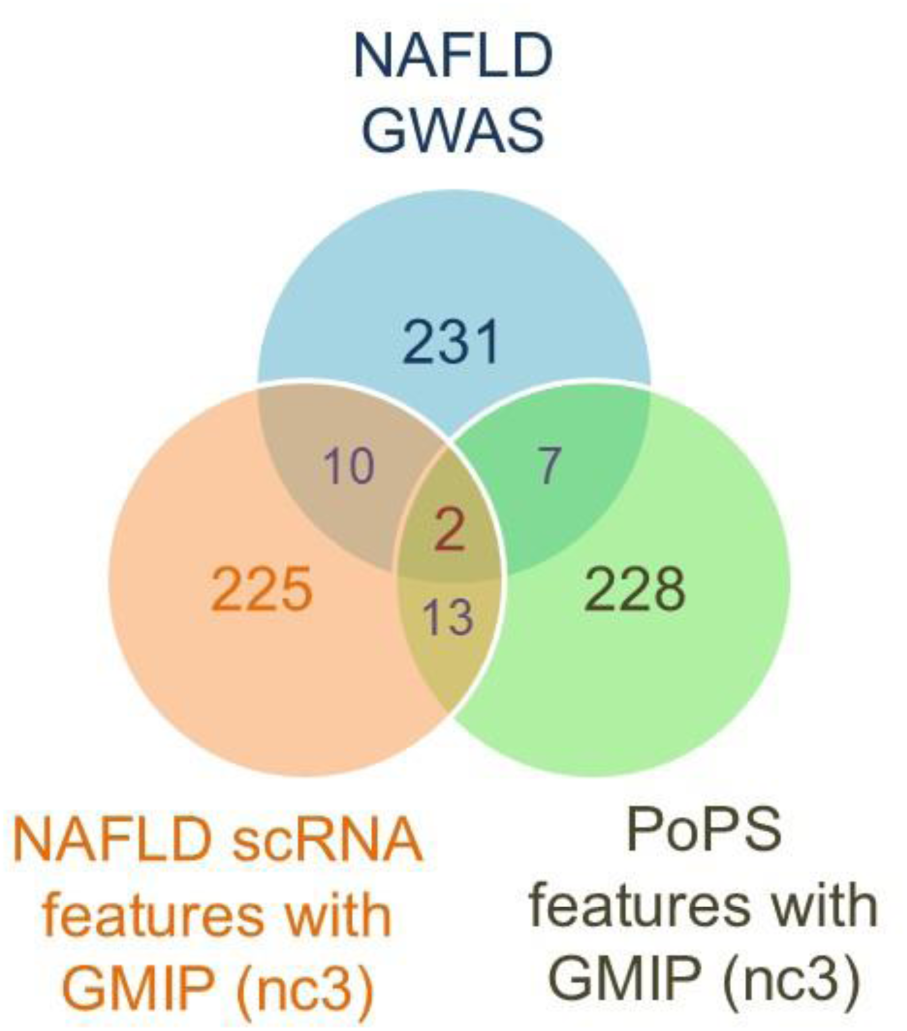
Overlap of NAFLD GWAS and Re-prioritization Gene Lists. Figure 7: Description We applied the GMIP pipeline to a NAFLD GWAS, utilizing both general PoPS features and specific features derived from a NAFLD mouse single-cell RNA-seq dataset. For the top 250 genes, the highest tau value of 2.96 was achieved using GMIP_PLSR_nc3 with PoPS features. When using scRNA-seq features and taking top 250 genes, the highest Tau value obtained was 1.59 also with GMIP_PLSR_nc3. Additionally, a third list of 250 genes was generated directly from the NAFLD MAGMA results. Also, the pairwise overlap of list is significant and as follows (Total genes=18383):

A. GWAS genes vs scRNA features genes: 0.0001687
B. GWAS genes vs PoPS features genes: 0.007394
C. scRNA features genes vs PoPS features genes: 1.798e-06 The Venn diagram shows the overlap between these gene lists. The marginal overlap indicates that each list captures different aspects of the data, highlighting the unique contributions of prioritization methods.

To further assess the functional relevance of these gene sets, we conducted pathway enrichment analysis using the DisGeNET and WikiPathways databases (Figure 8). The GMIP_PLSR_nc3 model with PoPS features identified a broader range of NAFLD-associated pathways (24 pathways), reflecting the capacity of general features to capture diverse biological processes. In contrast, the scRNA-seq features were linked to fewer pathways (4), suggesting a more focused interpretation of NAFLD biology. The original GWAS-derived gene set captured 6 pathways related to NAFLD. This comparison underscores the complementary strengths of disease-specific and general features, with PoPS features offering more comprehensive pathway coverage (Figure 8).

**Figure 8:**
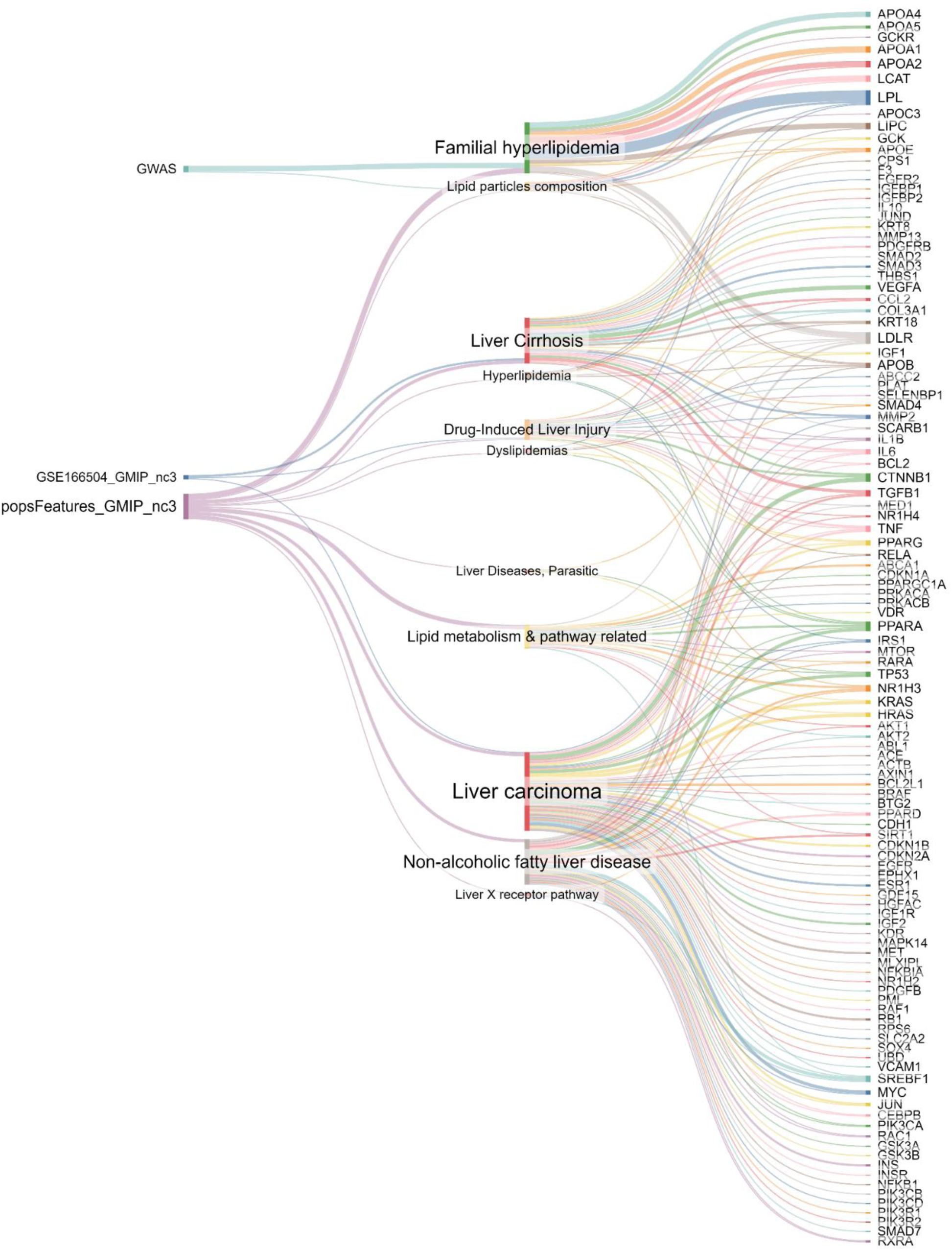
Pathway Enrichment Analysis of NAFLD Gene Lists. Figure 8: Description This Sankey plot visualizes the results of pathway enrichment analysis for the top 250 genes from three different top gene lists as shown in Figure 3.7. Over-Representation Analysis (ORA) was performed using DisGeNET and WikiPathways databases, focusing on pathways with an FDR < 0.1 that include the keywords “lipid” or “liver.“ The plot highlights the connections between the gene lists (left), the enriched pathways (middle), and the specific genes involved (right): **Gene List from PoPS Features with GMIP_PLSR_nc3**: This list was significantly associated with multiple pathways, including the “Non-alcoholic fatty liver disease” pathway from WikiPathways, indicating a strong link to NAFLD-related biological processes. **Original GWAS Gene List**: Primarily enriched for pathways related to “Familial hyperlipidemia,” as indicated by the flow. **Gene List from scRNA-seq Features with GMIP_PLSR_nc3**: Enriched for pathways related to “Drug-induced liver injury”, “Liver cirrhosis” and “Liver carcinoma,” suggesting a broader involvement in liver-related diseases. This is depicted by the connections from “GSE166504_GMIP_nc3” to relevant pathways.

## 4.0 Discussion

In this work, we developed and applied the GMIP (GWAS and Multi-omics Integration Pipeline) framework, integrating GWAS summary statistics with various multi-omics data. The flexible and modular design of GMIP allows for easy incorporation of widely used tools like PoPS, MAGMA, and NAGA, enabling gene reprioritization across a wide variety of GWAS. Furthermore, our implementation of GMIP-PLSR, which leverages Partial Least Squares Regression (PLSR) to address multicollinearity in PoPS-selected features, significantly enhanced gene prioritization in terms of both biological interpretability and computational efficiency.

### Strengths

#### Modularity and Flexibility

One of the key strengths of GMIP is its modular architecture, which allows users to select from a variety of methods and configurations for each analysis step. This flexibility enables comparative analyses and optimization across multiple datasets, making GMIP highly adaptable to different research scenarios. This is the first system of its kind, offering a unified and accessible platform for consistent analysis.

##### Multi-omics Integration

GMIP demonstrated its strength in integrating diverse omics data, including gene expression, protein-protein interactions, and curated pathways, to enhance the biological interpretation of GWAS results. By doing so, it provided a more holistic view of the genetic architecture underlying complex traits and diseases, as demonstrated in the NAFLD case study.

##### Multicollinearity Handling

By introducing GMIP-PLSR, we addressed a significant limitation in methods like PoPS, where multicollinearity in feature sets reduced the accuracy of gene ranking. PLSR efficiently mitigated this issue, improving both the stability and performance of the model, especially when applied to highly correlated multi-omics features. This will become even more important when more public data is added to the feature sets.

##### Scalability

Implemented using Nextflow (Di Tommaso et al., 2017), GMIP offers computational scalability, making it applicable in different environments from personal laptops to high-performance computing clusters. This adaptability is crucial for handling large-scale GWAS datasets and multi-omics data.

#### Weaknesses

##### Data Dependency

While GMIP excels in integrating multi-omics data, its performance heavily depends on the availability and quality of the underlying datasets. For example, in some cases, tissue-specific data or fine-grained single-cell datasets may not be available, limiting the method’s ability to prioritize genes accurately (Stuart et al., 2019).

##### Complexity for Non-Experts

Although GMIP is flexible, its modularity can also be a drawback for users unfamiliar with bioinformatics pipelines. Selecting the appropriate parameters, understanding the tools within each module, and interpreting the results require a certain level of expertise in computational genomics, which might pose challenges to researchers with limited experience in the field (Wilkinson et al., 2016).

##### Locus-based Methods

One area where GMIP can be further strengthened is in the combination of locus-based approaches. While tools like PoPS capture gene-level associations, further fine mapping or integrating locus-specific results, such as those from promoter capture Hi-C, could enhance the discovery of causal variants, offering even deeper insights into regulatory mechanisms (Javierre et al., 2016b).

### 5.0 Conclusion

This work presents GMIP as a comprehensive and scalable solution for gene prioritization by integrating GWAS and multi-omics data. By applying GMIP across multiple GWAS studies and demonstrating its effectiveness with PLSR integration, we provided a flexible framework that tackles critical challenges in post-GWAS analysis, such as multicollinearity and feature selection. GMIP has proven its utility in enhancing biological insights, particularly in complex diseases like NAFLD, where multi-omics data played a crucial role in gene reprioritization.

### 6.0 Perspectives

Looking ahead, several exciting avenues can further elevate the potential of GMIP for gene prioritization:

#### Combining Locus-Based Methods

One promising direction is to incorporate more locus-based fine-mapping methods like FINEMAP (Benner et al., 2016), PAINTOR (Kichaev et al., 2014), and promoter capture Hi-C alongside GMIP. This would allow for even more precise identification of causal variants by leveraging both gene- and locus-level signals. PoPS’s integration of locus-based methods could be a model for how GMIP could evolve in this direction.

#### Exploration of NAGA Features in GMIP-PLSR

The use of NAGA’s network features could be further explored in the GMIP-PLSR context. For instance, employing NAGA features alongside Partial Least Squares Regression could help further refine gene prioritization by capturing indirect network effects, thereby identifying novel disease-related pathways (Ou et al., 2024).

#### RNA-seq Foundational Models

With the emergence of foundational models (Hao et al., 2024) in RNA-seq, there is an exciting opportunity to use the latent features from these models as inputs into GMIP-PLSR. These features could represent a new dimension of gene regulation that is currently underutilized. By passing RNA-seq data through these models and integrating the derived features, GMIP could further improve gene ranking and interpretation in a wide range of complex traits.

#### Drug Discovery Pipelines

The modular nature of GMIP also makes it suitable for integration into drug discovery pipelines. By adding a drug prioritization module, GMIP could be applied to identify druggable targets, further extending its applicability to translational research (Han et al., 2021; Lotfi Shahreza et al., 2018). Additionally, exploring its use in combination with other pipelines like GPrioir, Priority Index, etc., which work differently and can give complementary results will be crucial.

## 8.0 Supplementary Tables and Figures

**Table S1:** Information on various GWAS studies used for evaluation. (Significance at 5e-8)

| TraitID | TraitName | N (sample size) | #SigSNPs |
| --- | --- | --- | --- |
| BMI | Body Mass Index | 361,194 | 49,224 |
| ECZ | Eczema | 458,699 | 17,361 |
| HDL | High Density Lipoprotein | 361,194 | 49,970 |
| LDL | Low Density Lipoprotein | 361,194 | 31,699 |
| NFD | Non-Alcoholic Fatty Liver Disease | 137,048 | 5,212 |
| RAD | Rheumatoid Arthritis | 57,284 | 20,686 |
| SCZ | Schizophrenia | 77,096 | 13,474 |
| T2D | Type 2 Diabetes | 459,324 | 3,250 |

**Table S2:**
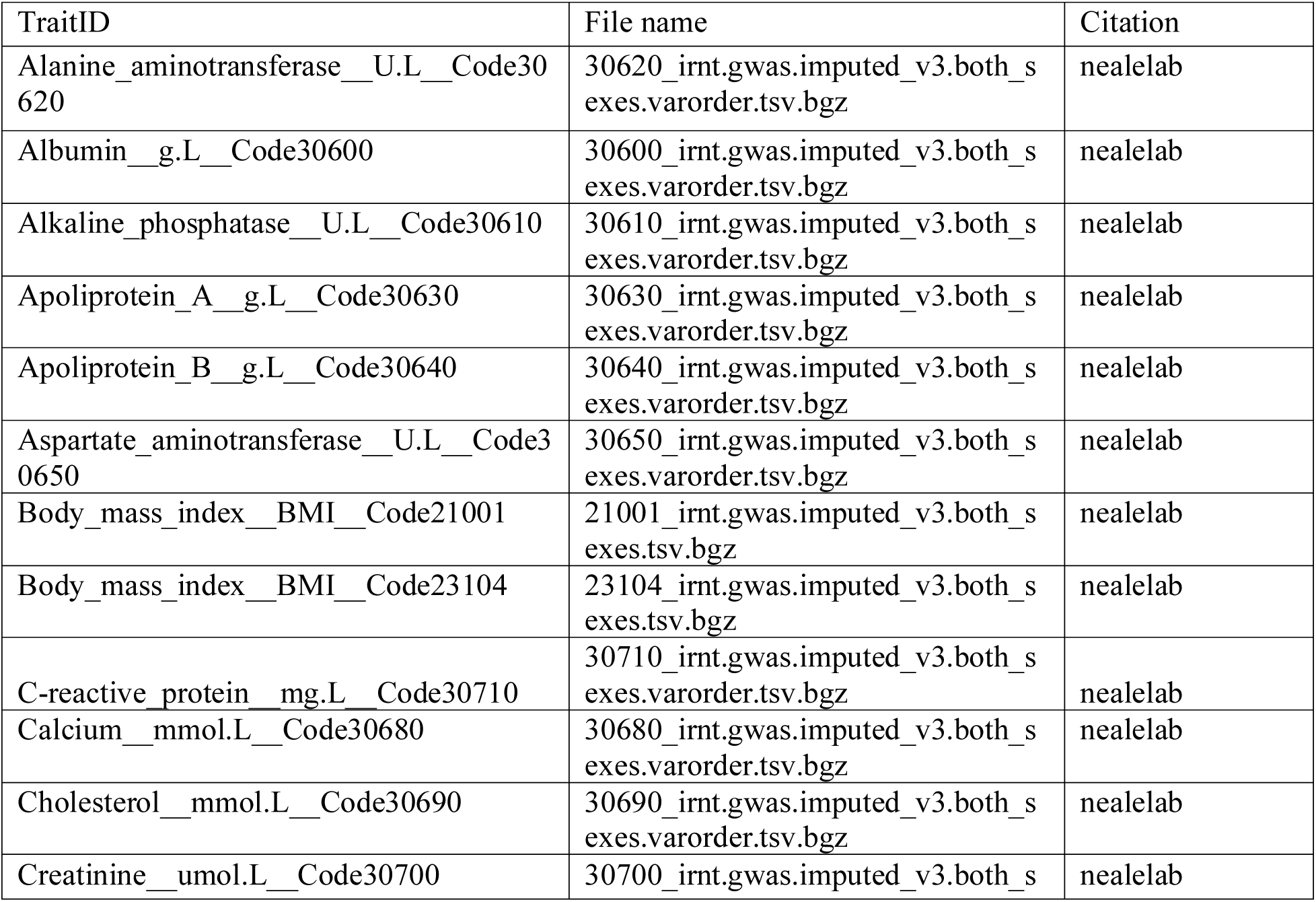
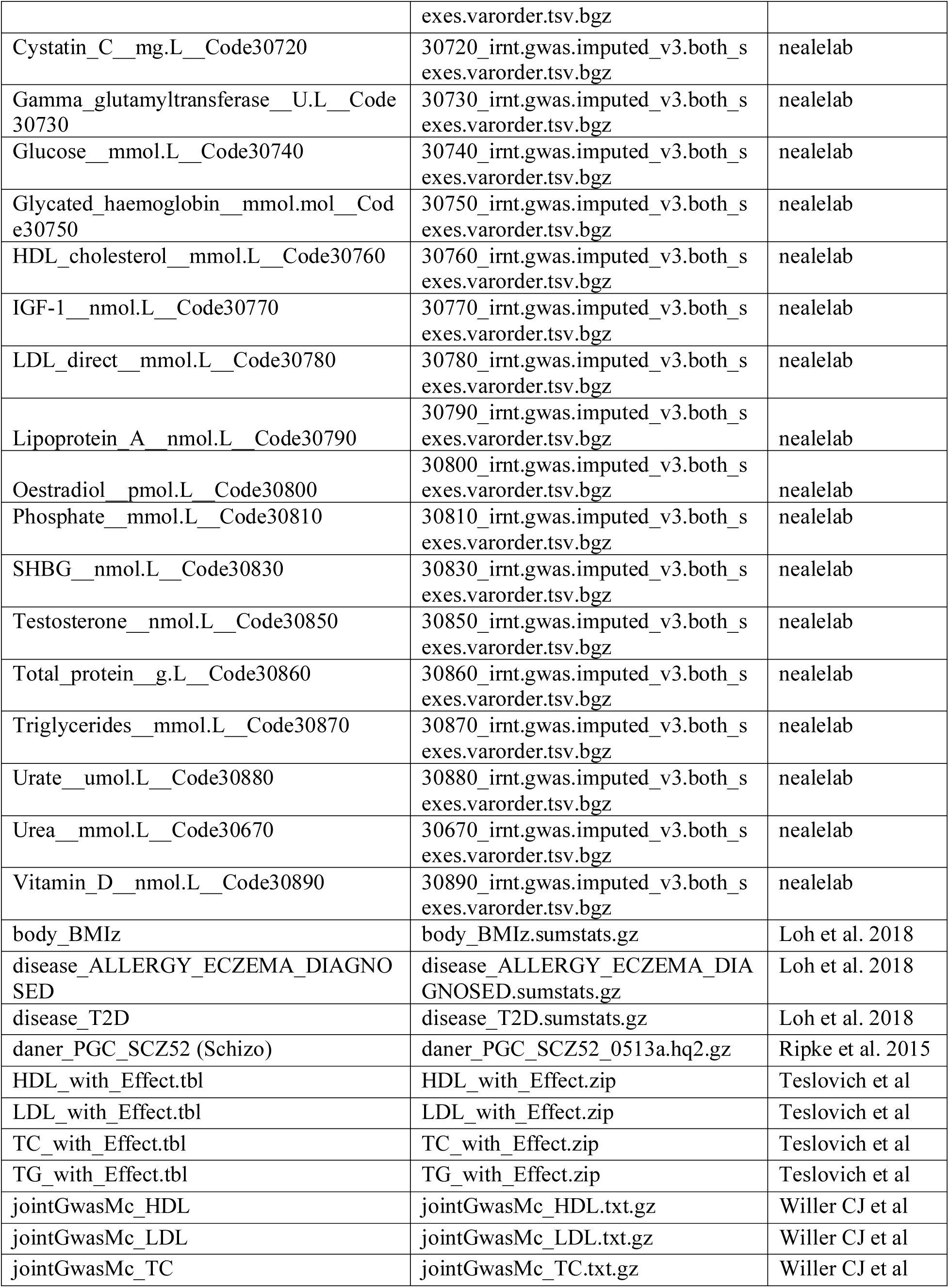
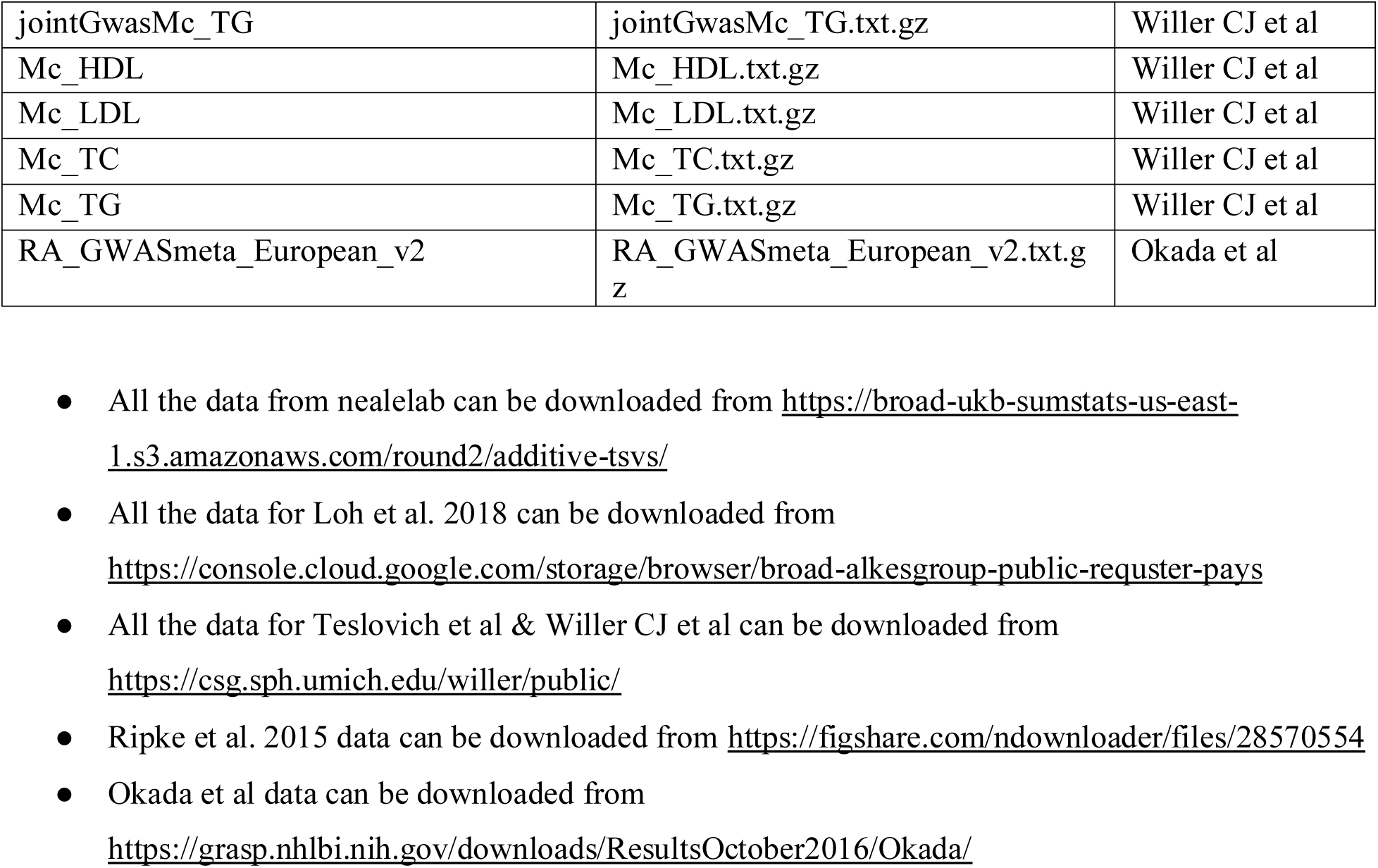
GWAS Source Details.

**Table S3:**
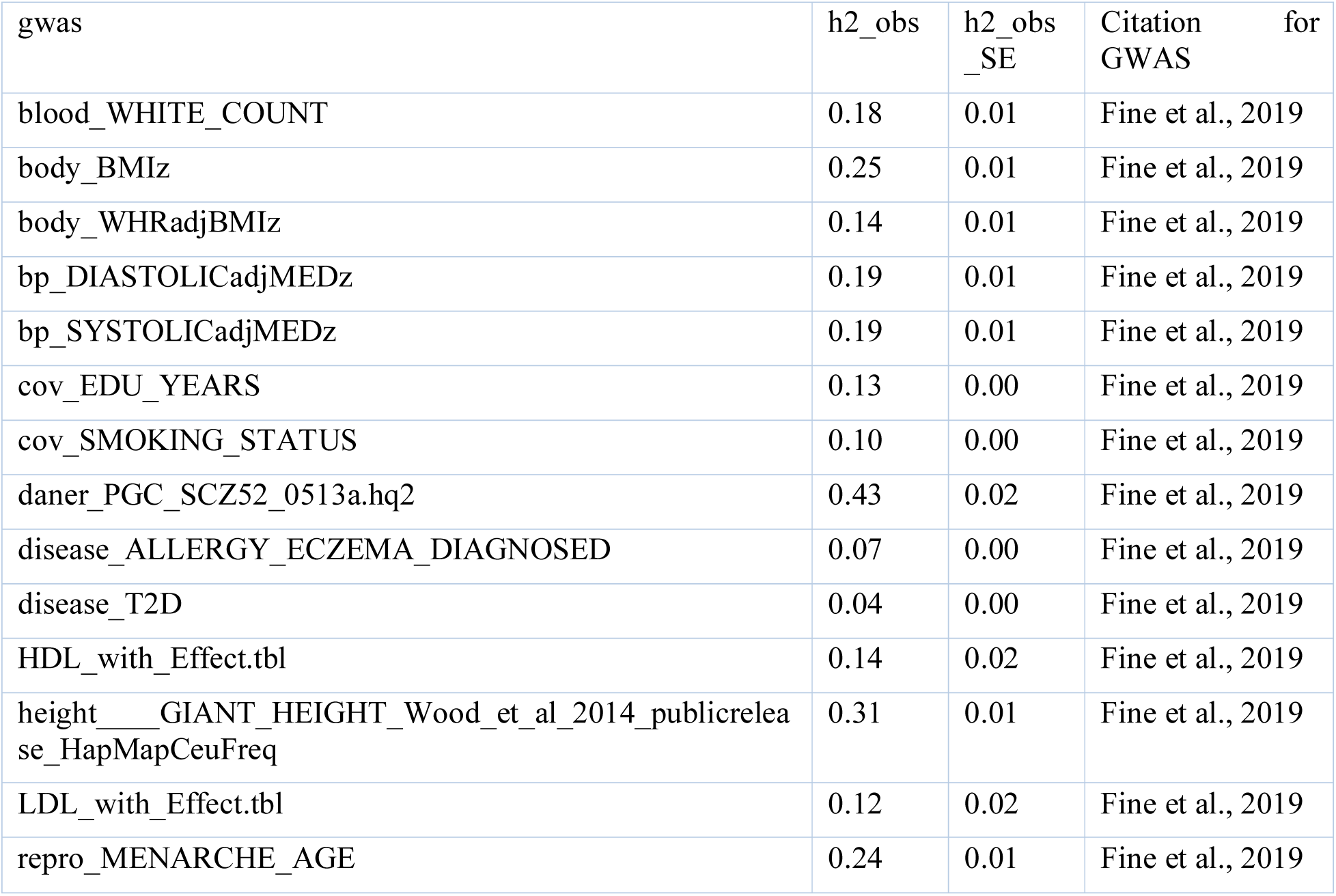
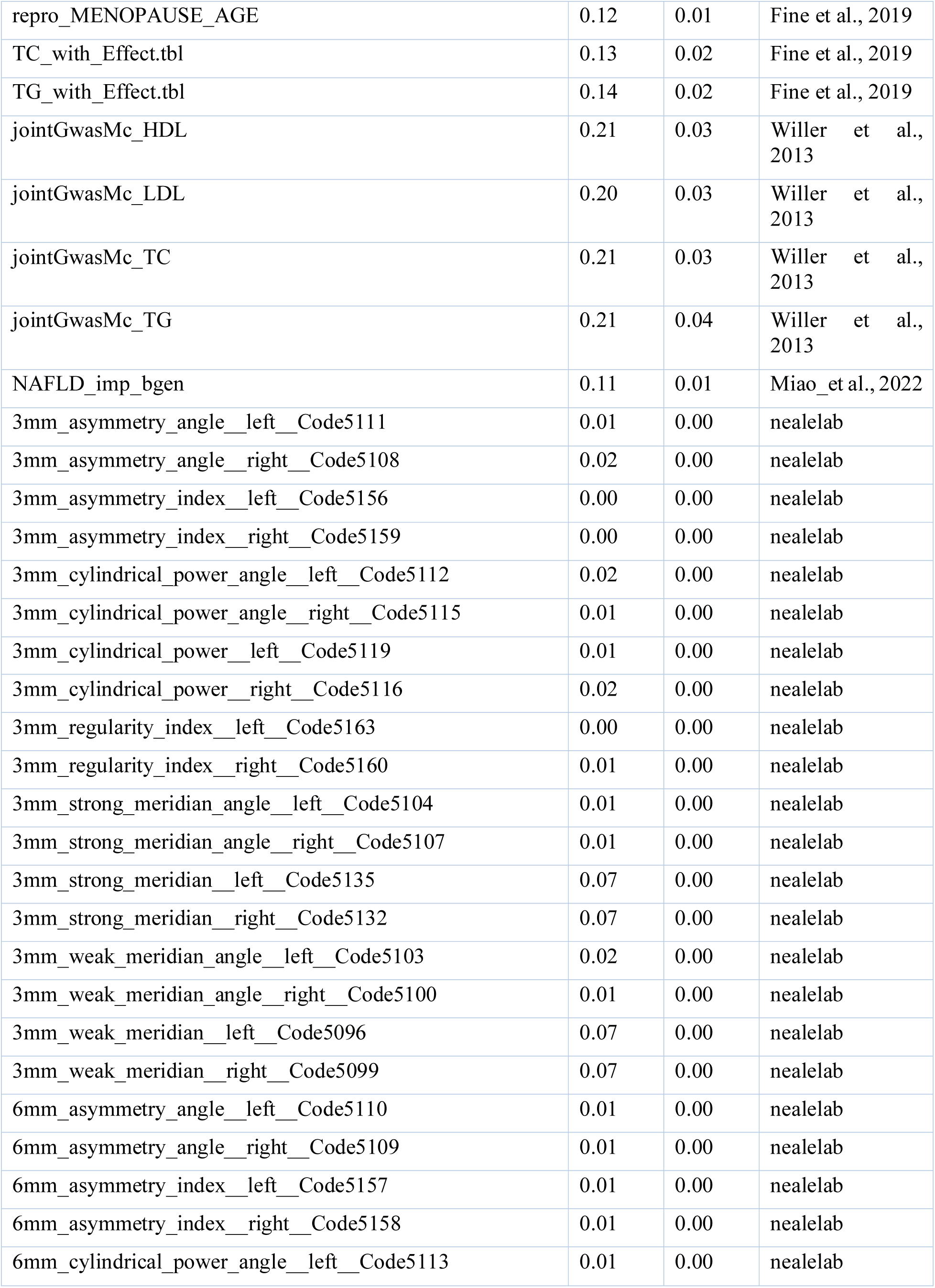
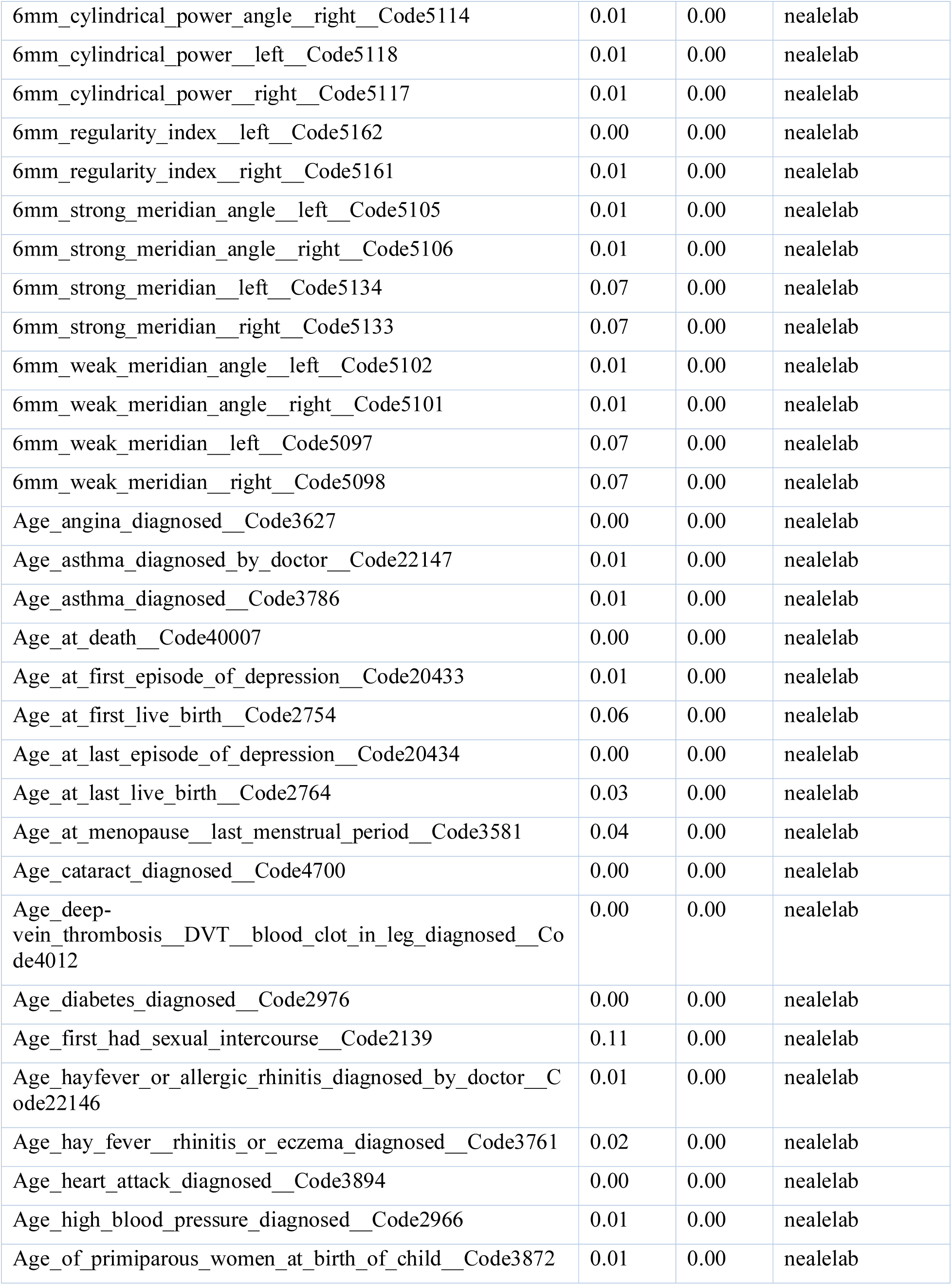
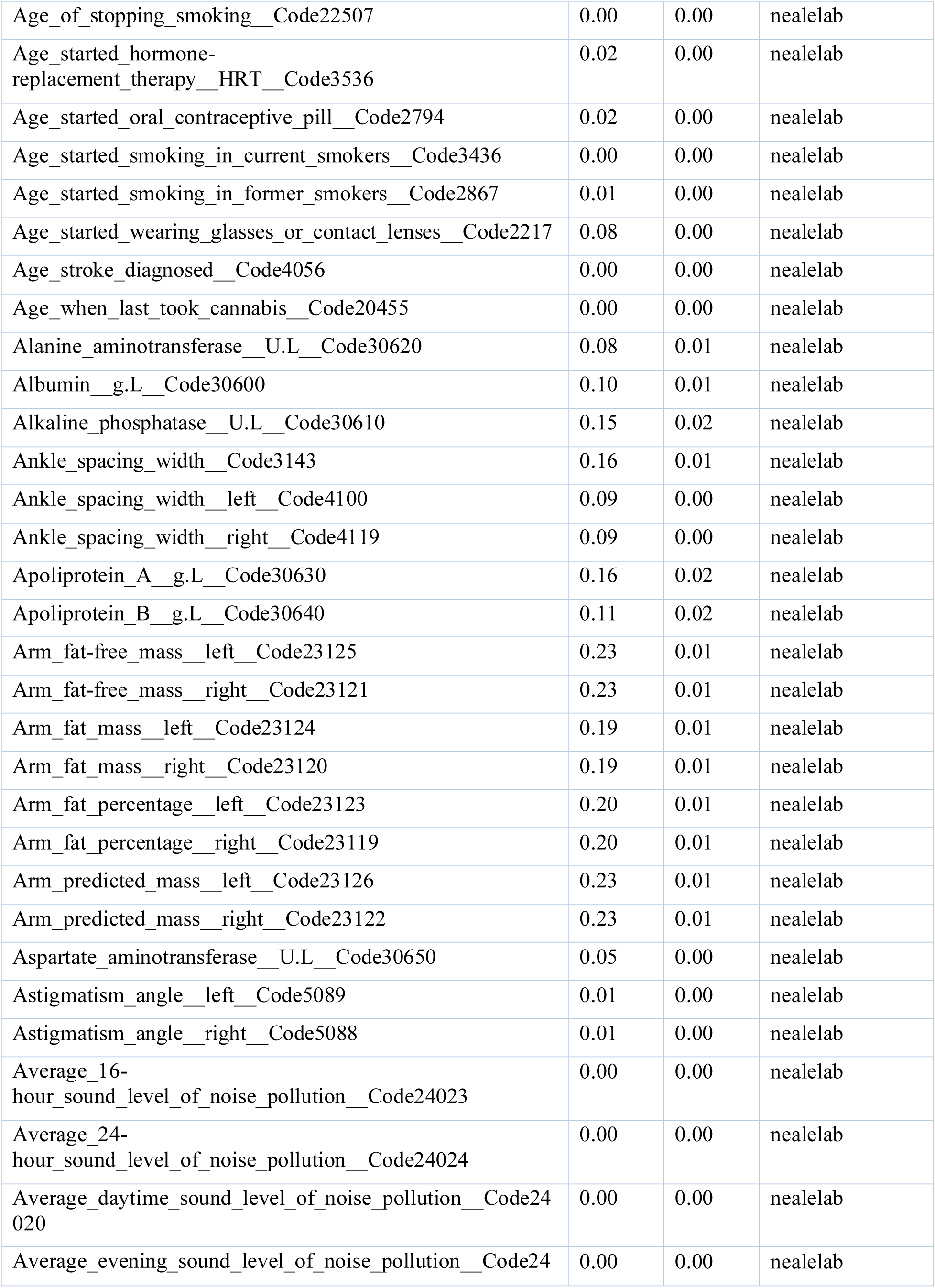
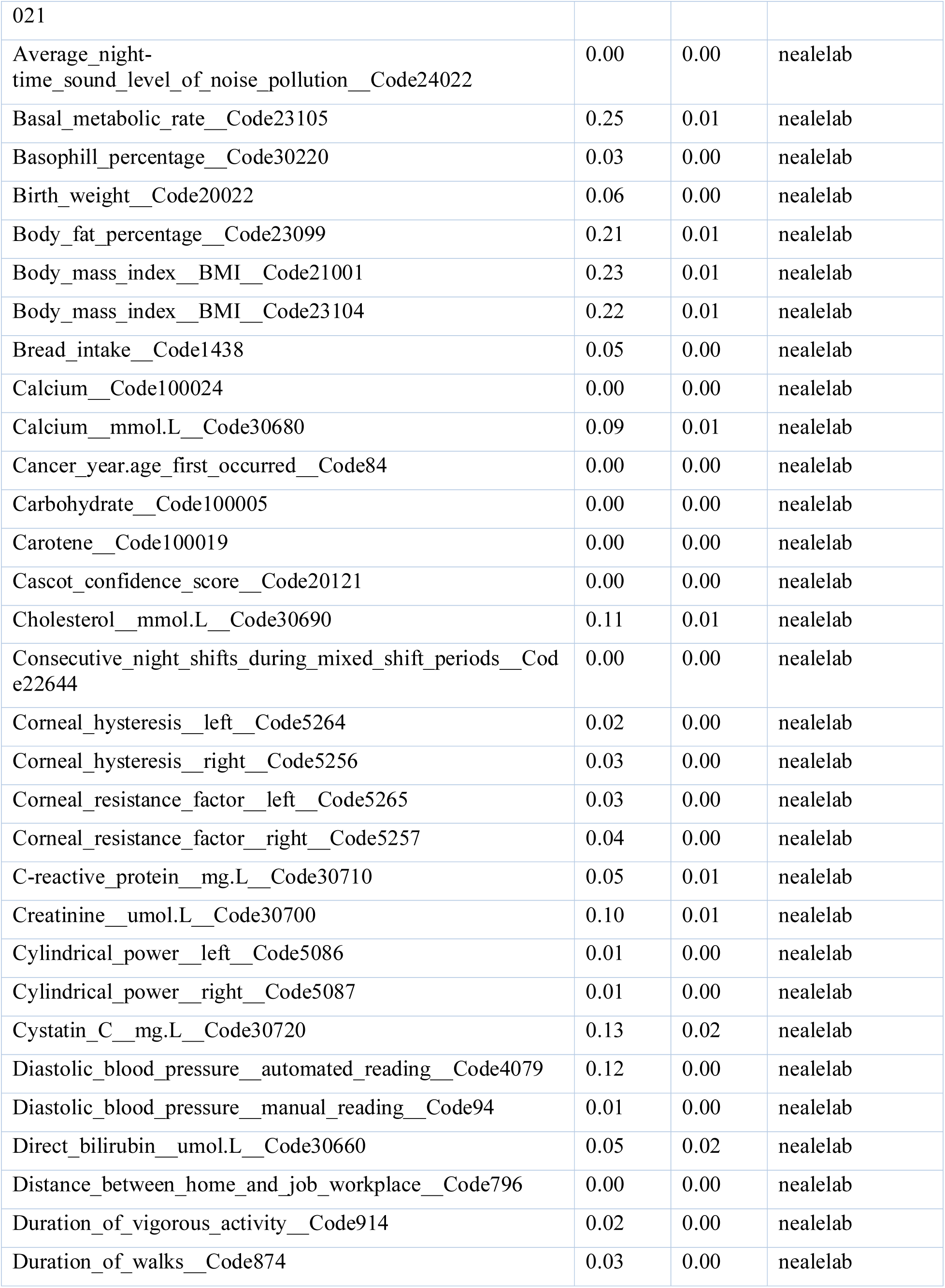
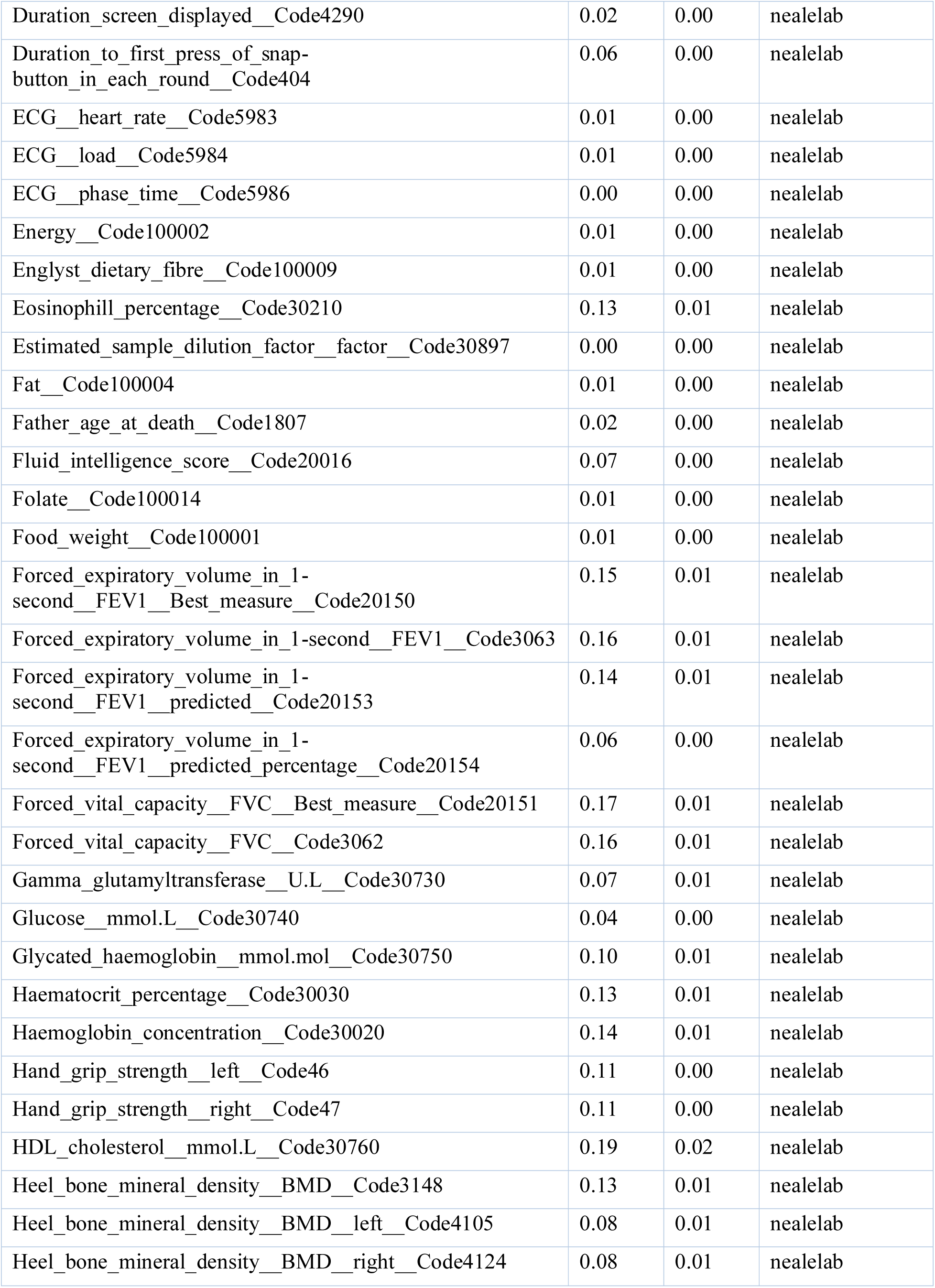
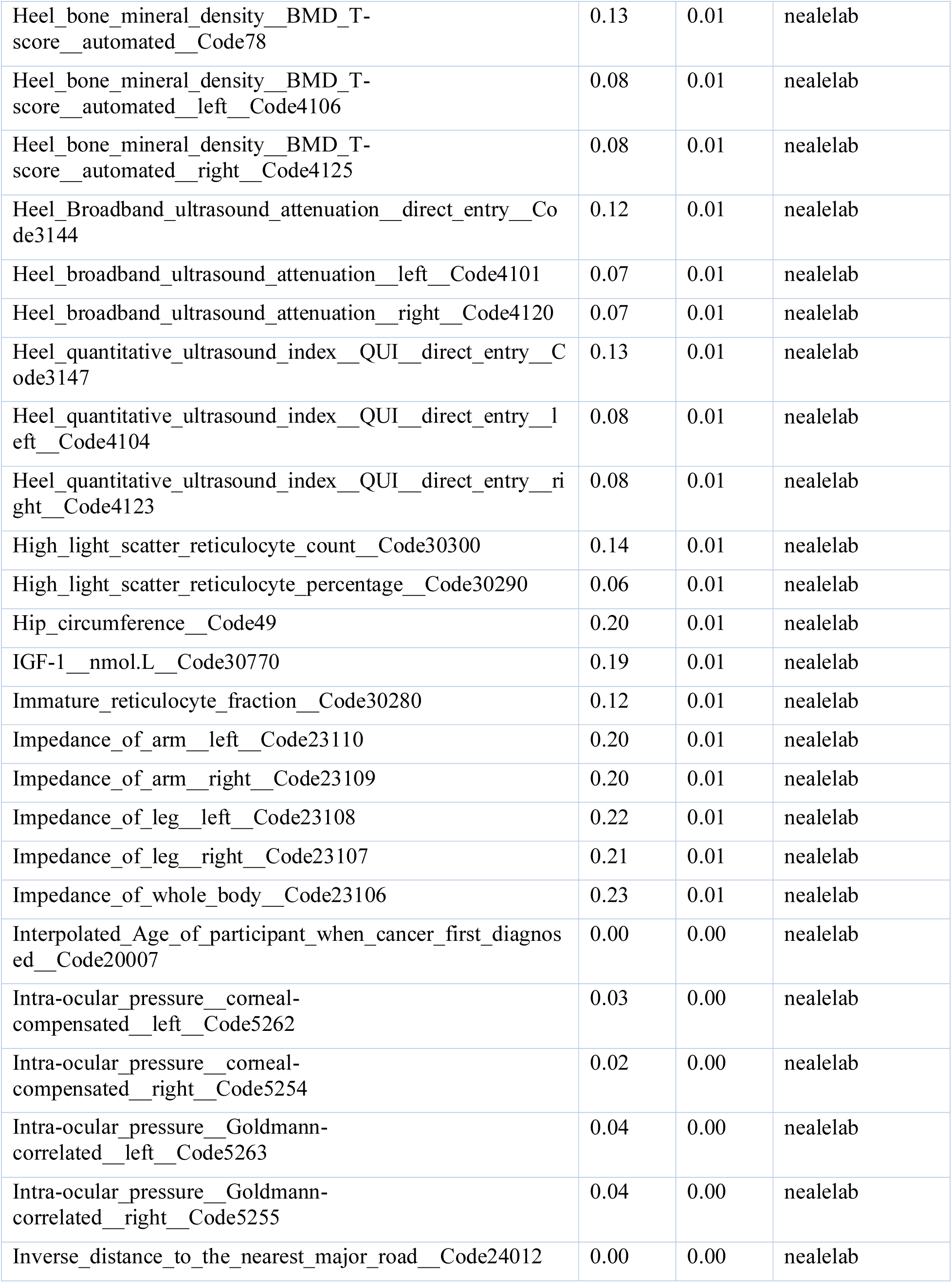
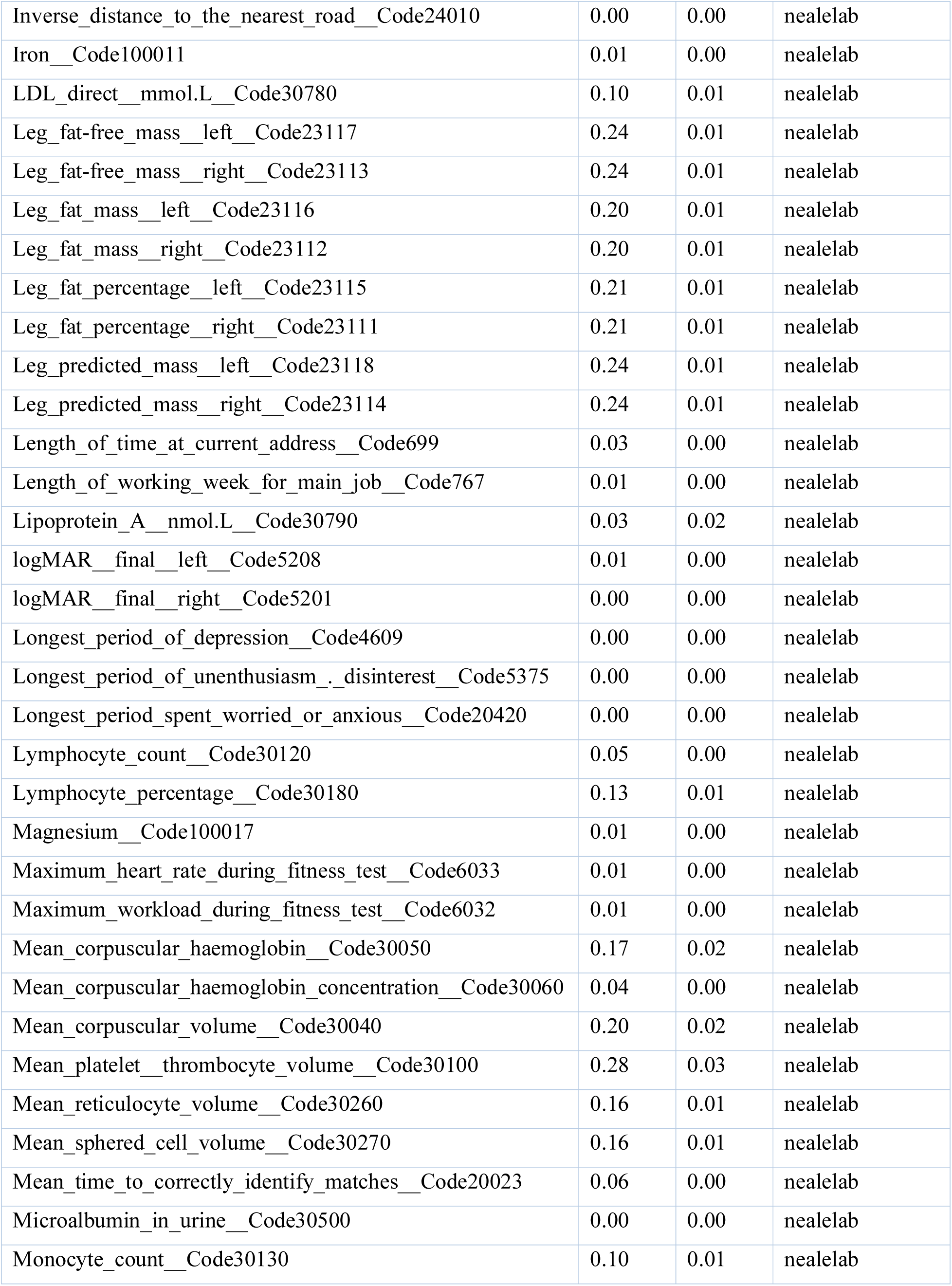
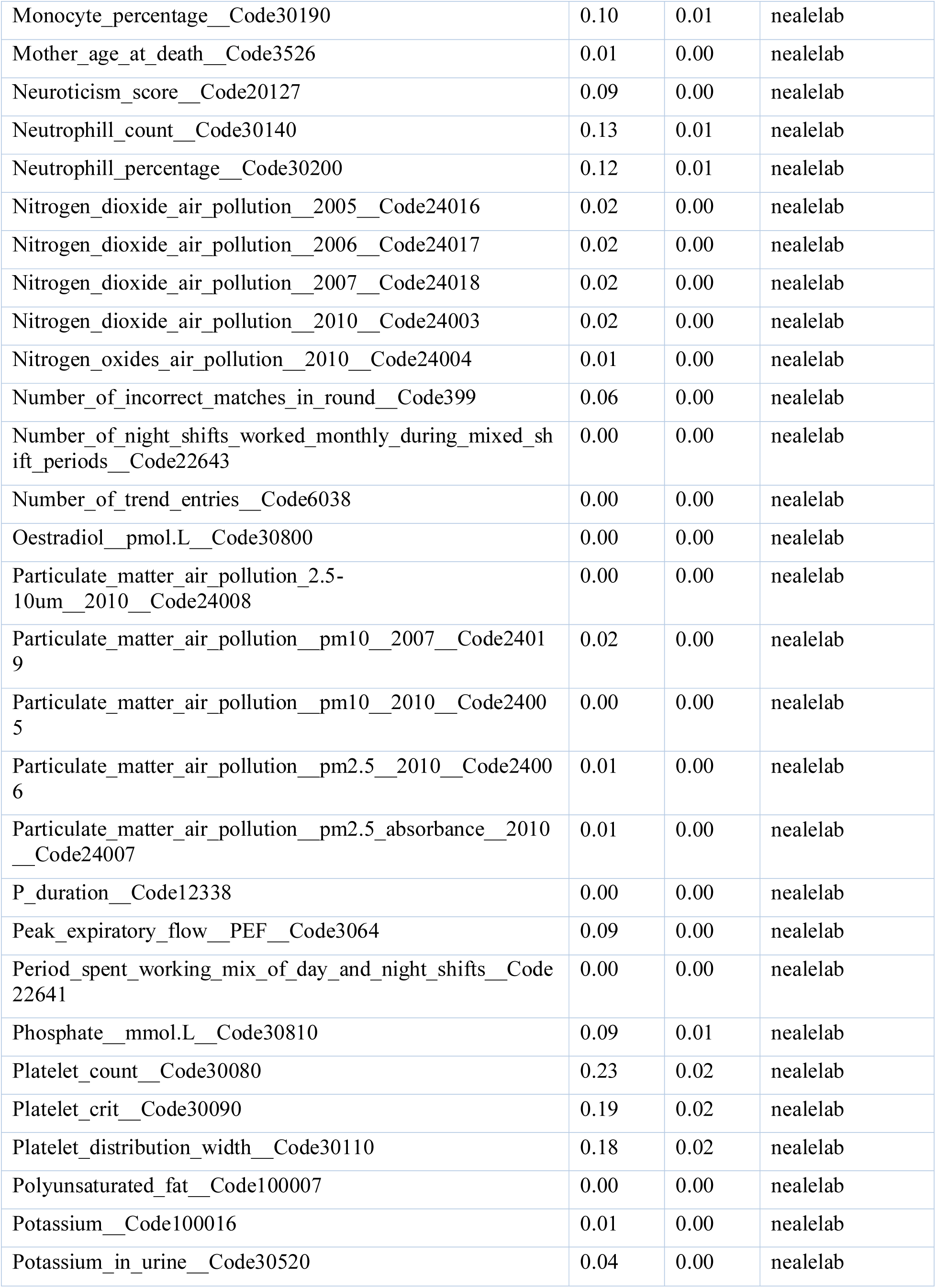
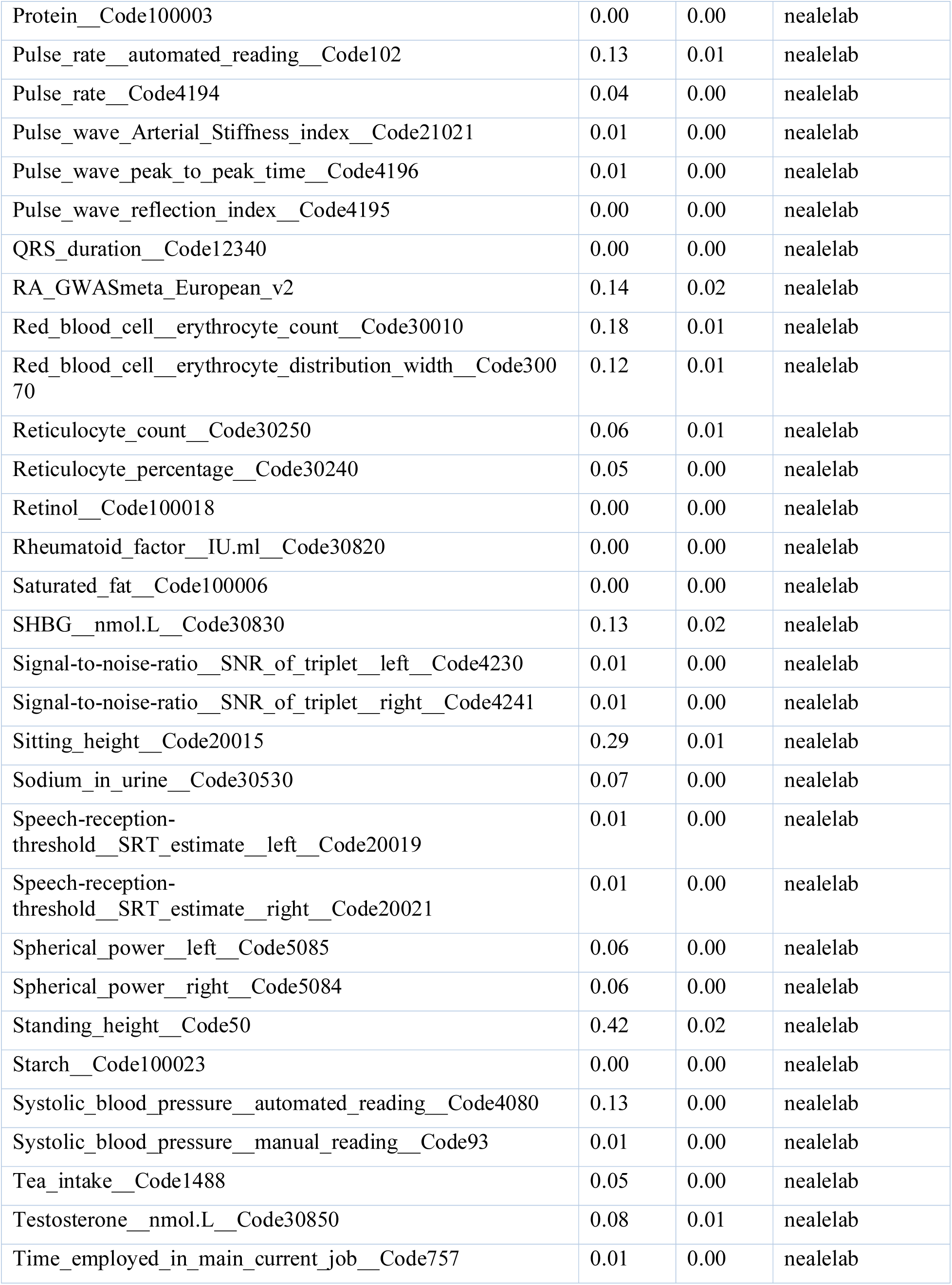
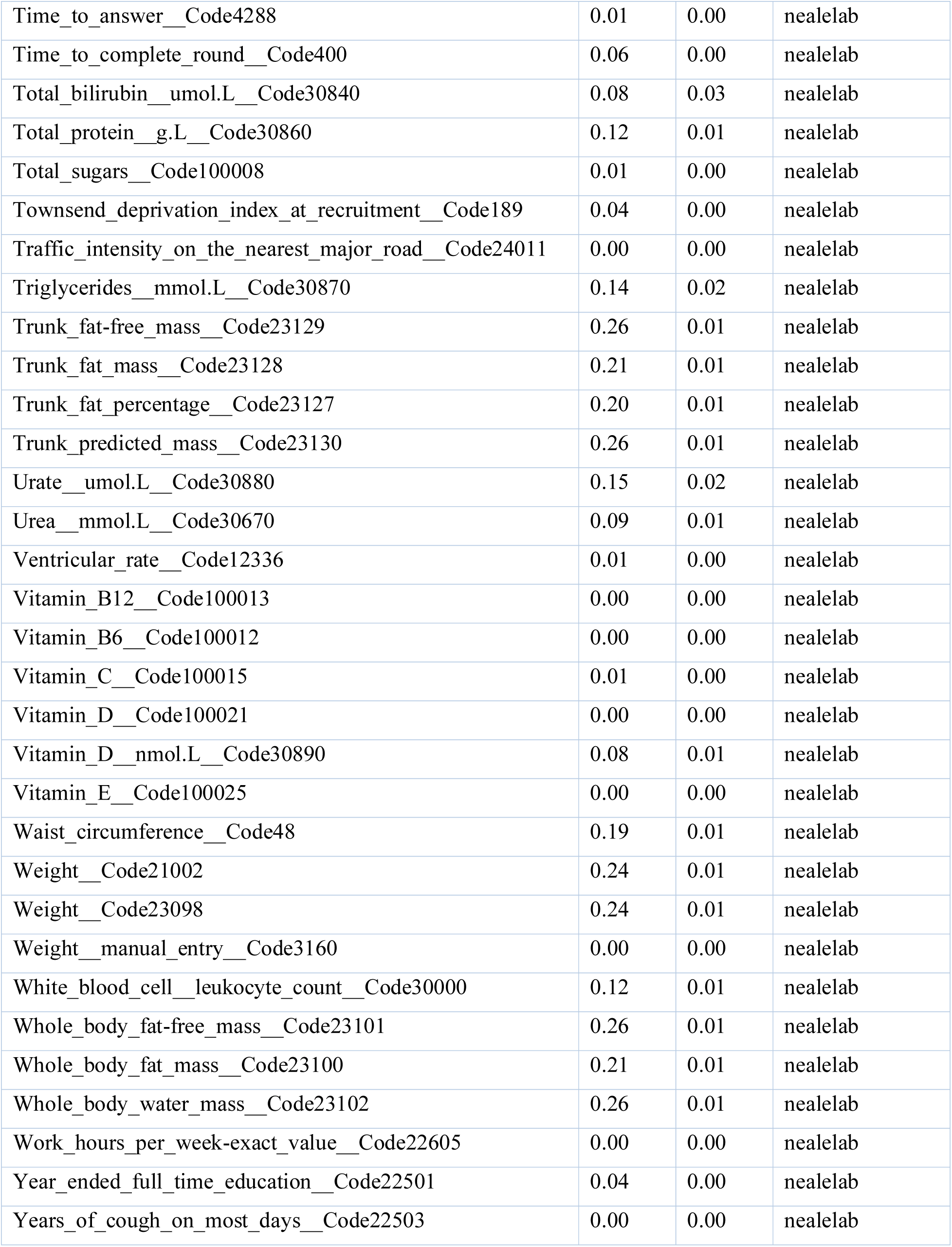
Observed Heritability calculated using LDSC.

**Figure S1:**
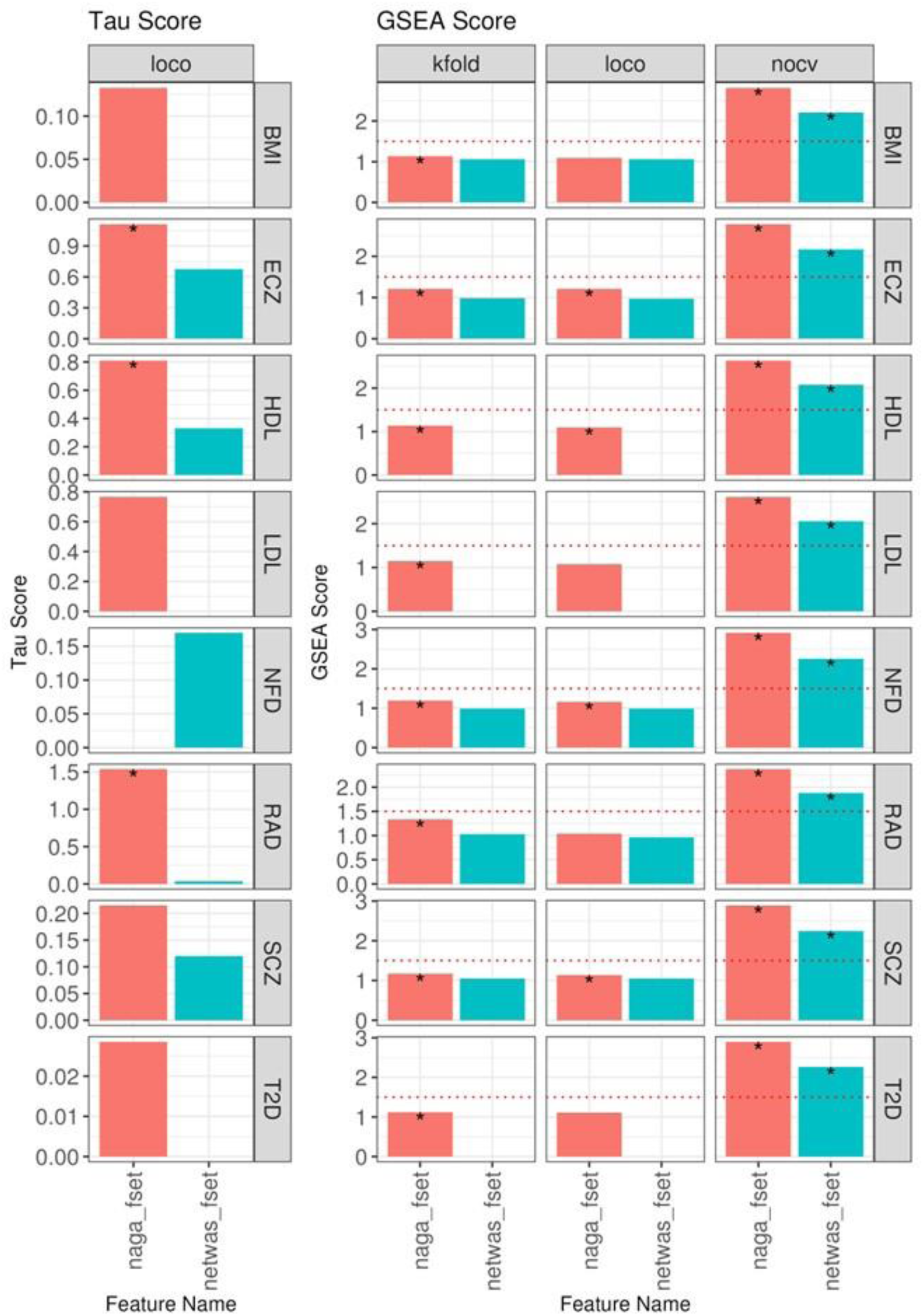
Performance of NAGA within GMIP. Figure S1: Description This figure illustrates the results of applying the NAGA method in GMIP, showing the comparative performance of tau and GSEA scores across eight different GWAS traits.

- **Left Panel**: Tau scores for the top 500 genes predicted by NAGA, evaluated using the LOCO cross-validation strategy for each GWAS trait.
- **Right Panel**: GSEA scores for three cross-validation strategies—k-fold, LOCO, and noCV. The gene set consists of the top 500 genes identified by MAGMA from the original GWAS, and the ranked list contains genes predicted by NAGA.
- **Rows:** Each row corresponds to a different GWAS trait (BMI, ECZ, T2D, etc.), comparing results between the NAGA-specific feature set (red) and the NetWAS feature set (blue).
- **Highlights:** Stars Mark significant p-values (<0.05), and red dotted lines represent the GSEA significance threshold (y = 1.5).
- Key Insights:

- **No Cross-Validation (noCV)**: NAGA feature sets performed well without cross-validation, yielding significant GSEA scores across most traits.
- **LOCO and k-fold Cross-Validation**: Both strategies often led to a reduction in performance, with lower GSEA and tau scores across many traits. However, NAGA showed better performance for the RAD trait. These results suggest that while NAGA’s feature sets perform well without cross-validation, the models tend to suffer from overfitting under LOCO and k-fold cross-validation, a common challenge with network propagation models.

**Figure S2:**
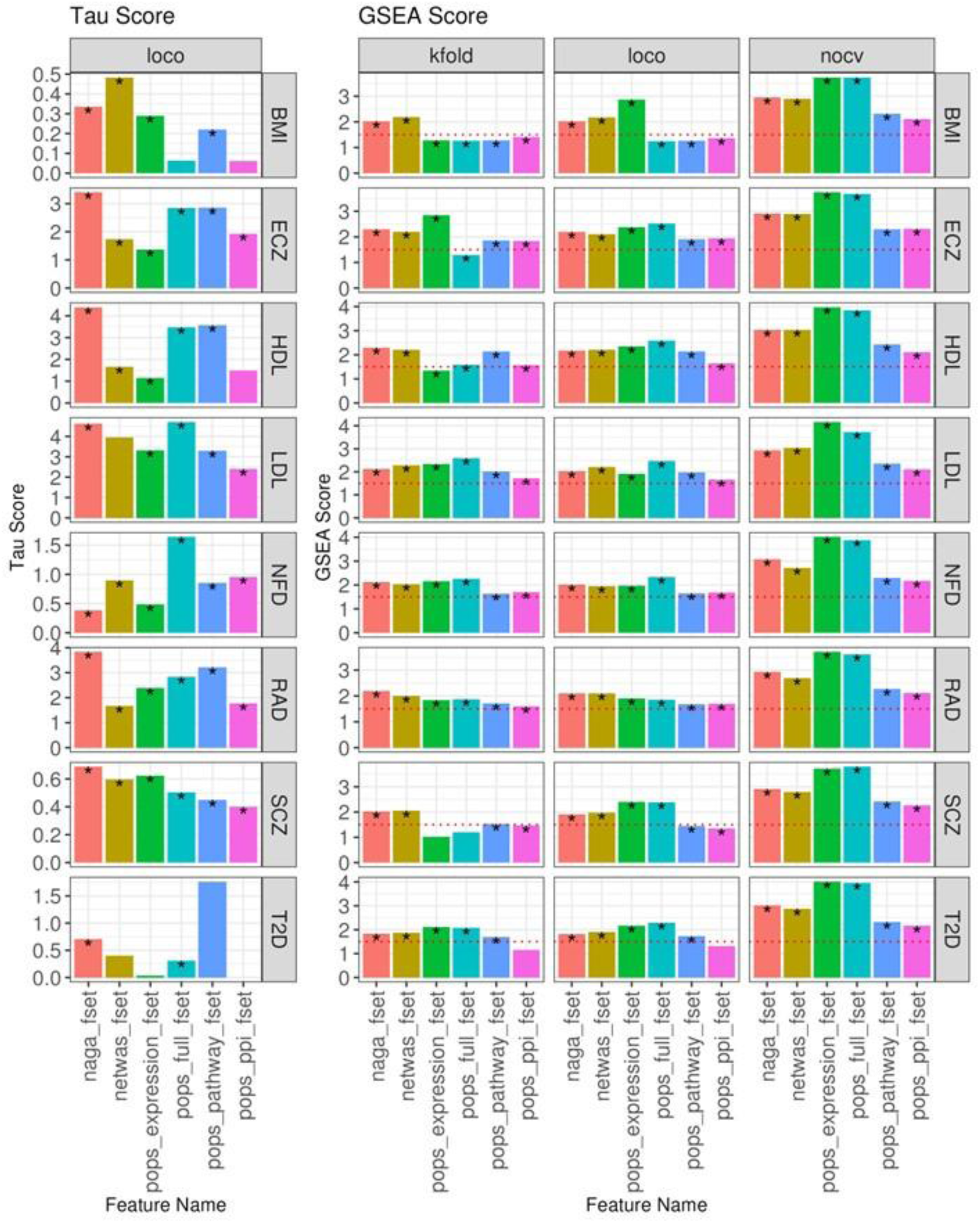
Performance of PoPS in GMIP. Figure S2 Description: This figure shows the performance of PoPS in GMIP, using various feature sets across eight GWAS traits.

- **Left Panel**: Tau scores for the top 500 genes predicted by PoPS using the LOCO cross-validation strategy.
- **Right Panel**: GSEA scores across k-fold, LOCO, and noCV cross-validation strategies. The top 500 genes from MAGMA z-scores are used as the gene set, and genes are ranked by PoPS predicted scores.
- **Rows**: Each row represents a GWAS trait, such as BMI, HDL, and T2D, comparing multiple feature sets like NAGA (PCnet), PoPS full, PoPS pathway, and PoPS expression.
- **Highlights:** Stars Mark significant p-values (<0.05), and red dotted lines represent the GSEA significance threshold (y = 1.5).
- Key Insights:

- PoPS method performs well across all trait and feature combinations
- The performance remains robust across various CV strategies
- NAGA features are as good or sometimes better for GWAS re-prioritization with PoPS This figure highlights the superior performance of PoPS across multiple GWAS traits and feature sets, demonstrating its generalizability and robustness.

## References

Auton, A., Abecasis, G. R., Altshuler, D. M., Durbin, R. M., Bentley, D. R., Chakravarti, A., Clark, A. G., Donnelly, P., Eichler, E. E., Flicek, P., Gabriel, S. B., Gibbs, R. A., Green, E. D., Hurles, M. E., Knoppers, B. M., Korbel, J. O., Lander, E. S., Lee, C., Lehrach, H., … Schloss, J. A. (2015). A global reference for human genetic variation. In Nature (Vol. 526, Issue 7571, pp. 68–74). Nature Publishing Group. 10.1038/nature15393

Belsley, D. A., Kuh, E., & Welsch, R. E. (1980). Regression Diagnostics: Identifying Influential Data and Sources of Collinearity. John Wiley & Sons.

Benner, C., Spencer, C. C. A., Havulinna, A. S., Salomaa, V., Ripatti, S., & Pirinen, M. (2016). FINEMAP: Efficient variable selection using summary data from genome-wide association studies. Bioinformatics, 32(10), 1493–1501. 10.1093/bioinformatics/btw018

Boulesteix, A. L., & Strimmer, K. (2007). Partial least squares: A versatile tool for the analysis of high-dimensional genomic data. Briefings in Bioinformatics, 8(1), 32–44. 10.1093/bib/bbl016

Buniello, A., Macarthur, J. A. L., Cerezo, M., Harris, L. W., Hayhurst, J., Malangone, C., McMahon, A., Morales, J., Mountjoy, E., Sollis, E., Suveges, D., Vrousgou, O., Whetzel, P. L., Amode, R., Guillen, J. A., Riat, H. S., Trevanion, S. J., Hall, P., Junkins, H., … Parkinson, H. (2019a). The NHGRI-EBI GWAS Catalog of published genome-wide association studies, targeted arrays and summary statistics 2019. Nucleic Acids Research, 47(D1), D1005–D1012. 10.1093/nar/gky1120

Carlin, D. E., Fong, S. H., Qin, Y., Jia, T., Huang, J. K., Bao, B., Zhang, C., & Ideker, T. (2019). A Fast and Flexible Framework for Network-Assisted Genomic Association. IScience, 16, 155–161. 10.1016/j.isci.2019.05.025

Carrascal, L. M., Galván, I., & Gordo, O. (2009). Partial least squares regression as an alternative to current regression methods used in ecology. Oikos, 118(5), 681–690. 10.1111/j.1600-0706.2008.16881.x

Cowen, L., Ideker, T., Raphael, B. J., & Sharan, R. (2017). Network propagation: A universal amplifier of genetic associations. In Nature Reviews Genetics (Vol. 18, Issue 9, pp. 551–562). Nature Publishing Group. 10.1038/nrg.2017.38

de Leeuw, C. A., Mooij, J. M., Heskes, T., & Posthuma, D. (2015). MAGMA: Generalized Gene-Set Analysis of GWAS Data. PLoS Computational Biology, 11(4). 10.1371/journal.pcbi.1004219

Di Tommaso, P., Chatzou, M., Floden, E. W., Barja, P. P., Palumbo, E., & Notredame, C. (2017). Nextflow enables reproducible computational workflows. Nature Biotechnology, 35(4), 316–319. 10.1038/nbt.3820

Dietterich, T. G. (1998). Approximate Statistical Tests for Comparing Supervised Classification Learning Algorithms. Neural Computation, 10(7), 1895–1923. 10.1162/089976698300017197

Fine, R. S., Pers, T. H., Amariuta, T., Raychaudhuri, S., & Hirschhorn, J. N. (2019). Benchmarker: An Unbiased, Association-Data-Driven Strategy to Evaluate Gene Prioritization Algorithms. American Journal of Human Genetics, 104(6), 1025–1039. 10.1016/j.ajhg.2019.03.027

Frank, E., & Friedman, J. H. (1993). A Statistical View of Some Chemometrics Regression Tools. In Control TECHNOMETRICS (Vol. 35, Issue 2).

Gamazon, E. R., Wheeler, H. E., Shah, K. P., Mozaffari, S. V., Aquino-Michaels, K., Carroll, R. J., Eyler, A. E., Denny, J. C., Nicolae, D. L., Cox, N. J., & Im, H. K. (2015). A gene-based association method for mapping traits using reference transcriptome data. Nature Genetics, 47(9), 1091–1098. 10.1038/ng.3367

Grant, C. E., Bailey, T. L., & Noble, W. S. (2011). FIMO: Scanning for occurrences of a given motif. Bioinformatics, 27(7), 1017–1018. 10.1093/bioinformatics/btr064

Greene, C. S., Krishnan, A., Wong, A. K., Ricciotti, E., Zelaya, R. A., Himmelstein, D. S., Zhang, R., Hartmann, B. M., Zaslavsky, E., Sealfon, S. C., Chasman, D. I., Fitzgerald, G. A., Dolinski, K., Grosser, T., & Troyanskaya, O. G. (2015). Understanding multicellular function and disease with human tissue-specific networks. Nature Genetics, 47(6), 569–576. 10.1038/ng.3259

Greene, W. H.. (2003). Econometric analysis : 5th ed. Prentice Hall.

Gusev, A., Ko, A., Shi, H., Bhatia, G., Chung, W., Penninx, B. W. J. H., Jansen, R., De Geus, E. J. C., Boomsma, D. I., Wright, F. A., Sullivan, P. F., Nikkola, E., Alvarez, M., Civelek, M., Lusis, A. J., Lehtimäki, T., Raitoharju, E., Kähönen, M., Seppälä, I., … Pasaniuc, B. (2016). Integrative approaches for large-scale transcriptome-wide association studies. Nature Genetics, 48(3), 245–252. 10.1038/ng.3506

Han, Y., Wang, C., Klinger, K., Rajpal, D. K., & Zhu, C. (2021). An integrative network-based approach for drug target indication expansion. PLoS ONE, 16(7 July). 10.1371/journal.pone.0253614

Hao, M., Gong, J., Zeng, X., Liu, C., Guo, Y., Cheng, X., Wang, T., Ma, J., Zhang, X., & Song, L. (2024). Large-scale foundation model on single-cell transcriptomics. Nature Methods, 21(8), 1481–1491. 10.1038/s41592-024-02305-7

Harrison, P. W., Amode, M. R., Austine-Orimoloye, O., Azov, A. G., Barba, M., Barnes, I., Becker, A., Bennett, R., Berry, A., Bhai, J., Bhurji, S. K., Boddu, S., Lins, P. R. B., Brooks, L., Ramaraju, S. B., Campbell, L. I., Martinez, M. C., Charkhchi, M., Chougule, K., … Yates, A. D. (2024). Ensembl 2024. Nucleic Acids Research, 52(D1), D891–D899. 10.1093/nar/gkad1049

Hasin, Y., Seldin, M., & Lusis, A. (2017). Multi-omics approaches to disease. In Genome Biology (Vol. 18, Issue 1). BioMed Central Ltd. 10.1186/s13059-017-1215-1

Hindorff, L. A., Sethupathy, P., Junkins, H. A., Ramos, E. M., Mehta, J. P., Collins, F. S., & Manolio, T. A. (n.d.). Potential etiologic and functional implications of genome-wide association loci for human diseases and traits.

Hormozdiari, F., van de Bunt, M., Segrè, A. V., Li, X., Joo, J. W. J., Bilow, M., Sul, J. H., Sankararaman, S., Pasaniuc, B., & Eskin, E. (2016). Colocalization of GWAS and eQTL Signals Detects Target Genes. American Journal of Human Genetics, 99(6), 1245–1260. 10.1016/j.ajhg.2016.10.003

Huang, J. K., Carlin, D. E., Yu, M. K., Zhang, W., Kreisberg, J. F., Tamayo, P., & Ideker, T. (2018). Systematic Evaluation of Molecular Networks for Discovery of Disease Genes. Cell Systems, 6(4), 484–495.e5. 10.1016/j.cels.2018.03.001

Javierre, B. M., Sewitz, S., Cairns, J., Wingett, S. W., Várnai, C., Thiecke, M. J., Freire-Pritchett, P., Spivakov, M., Fraser, P., Burren, O. S., Cutler, A. J., Todd, J. A., Wallace, C., Wilder, S. P., Kreuzhuber, R., Kostadima, M., Zerbino, D. R., Stegle, O., Burden, F., … Flicek, P. (2016a). Lineage-Specific Genome Architecture Links Enhancers and Non-coding Disease Variants to Target Gene Promoters. Cell, 167(5), 1369–1384.e19. 10.1016/j.cell.2016.09.037

Johnson, T. E., Lithgow, G. J., Murakami, S., & Shook, D. R. (2007). Genetics (2nd ed.). Elsevier Inc.

Kerrien, S., Aranda, B., Breuza, L., Bridge, A., Broackes-Carter, F., Chen, C., Duesbury, M., Dumousseau, M., Feuermann, M., Hinz, U., Jandrasits, C., Jimenez, R. C., Khadake, J., Mahadevan, U., Masson, P., Pedruzzi, I., Pfeiffenberger, E., Porras, P., Raghunath, A., … Hermjakob, H. (2012). The IntAct molecular interaction database in 2012. Nucleic Acids Research, 40(D1). 10.1093/nar/gkr1088

Kichaev, G., Yang, W. Y., Lindstrom, S., Hormozdiari, F., Eskin, E., Price, A. L., Kraft, P., & Pasaniuc, B. (2014). Integrating Functional Data to Prioritize Causal Variants in Statistical Fine-Mapping Studies. PLoS Genetics, 10(10). 10.1371/journal.pgen.1004722

Kolosov, N., Daly, M. J., & Artomov, M. (2021). Prioritization of disease genes from GWAS using ensemble-based positive-unlabeled learning. European Journal of Human Genetics, 29(10), 1527–1535. 10.1038/s41431-021-00930-w

Lander, S., Linton, L. M., Birren, B., Nusbaum, C., Zody, M. C., Baldwin, J., Devon, K., Dewar, K., Doyle, M., FitzHugh, W., Funke, R., Gage, D., Harris, K., Heaford, A., Howland, J., Kann, L., Lehoczky, J., LeVine, R., McEwan, P., … Yeh, R.-F. (2001). Initial sequencing and analysis of the human genome International Human Genome Sequencing Consortium* The Sanger Centre: Beijing Genomics Institute/Human Genome Center. In NATURE (Vol. 409). www.nature.com

Lê Cao, K. A., González, I., & Déjean, S. (2009). IntegrOmics: An R package to unravel relationships between two omics datasets. Bioinformatics, 25(21), 2855–2856. 10.1093/bioinformatics/btp515

Li, T., Wernersson, R., Hansen, R. B., Horn, H., Mercer, J., Slodkowicz, G., Workman, C. T., Rigina, O., Rapacki, K., Stærfeldt, H. H., Brunak, S., Jensen, T. S., & Lage, K. (2017). A scored human protein–protein interaction network to catalyze genomic interpretation. Nature Methods, 14(1), 61–64. 10.1038/nmeth.4083

Licata, L., Briganti, L., Peluso, D., Perfetto, L., Iannuccelli, M., Galeota, E., Sacco, F., Palma, A., Nardozza, A. P., Santonico, E., Castagnoli, L., & Cesareni, G. (2012). MINT, the molecular interaction database: 2012 Update. Nucleic Acids Research, 40(D1). 10.1093/nar/gkr930

Lonsdale, J., Thomas, J., Salvatore, M., Phillips, R., Lo, E., Shad, S., Hasz, R., Walters, G., Garcia, F., Young, N., Foster, B., Moser, M., Karasik, E., Gillard, B., Ramsey, K., Sullivan, S., Bridge, J., Magazine, H., Syron, J., … Moore, H. F. (2013). The Genotype-Tissue Expression (GTEx) project. In Nature Genetics (Vol. 45, Issue 6, pp. 580–585). 10.1038/ng.2653

Lotfi Shahreza, M., Ghadiri, N., Mousavi, S. R., Varshosaz, J., & Green, J. R. (2018). A review of network-based approaches to drug repositioning. In Briefings in bioinformatics (Vol. 19, Issue 5, pp. 878–892). NLM (Medline). 10.1093/bib/bbx017

Manolio, T. A. (2013). Bringing genome-wide association findings into clinical use. In Nature Reviews Genetics (Vol. 14, Issue 8, pp. 549–558). 10.1038/nrg3523

Marbach, D., Lamparter, D., Quon, G., Kellis, M., Kutalik, Z., & Bergmann, S. (2016). Tissue-specific regulatory circuits reveal variable modular perturbations across complex diseases. Nature Methods, 13(4), 366–370. 10.1038/nmeth.3799

Maurano, M. T., Humbert, R., Rynes, E., Thurman, R. E., Haugen, E., Wang, H., Reynolds, A. P., Sandstrom, R., Qu, H., Brody, J., Shafer, A., Neri, F., Lee, K., Kutyavin, T., Stehling-Sun, S., Johnson, A. K., Canfield, T. K., Giste, E., Diegel, M., … Stamatoyannopoulos, J. A. (n.d.). Systematic Localization of Common Disease-Associated Variation in Regulatory DNA. https://www.science.org

Mbatchou, J., Barnard, L., Backman, J., Marcketta, A., Kosmicki, J. A., Ziyatdinov, A., Benner, C., O’Dushlaine, C., Barber, M., Boutkov, B., Habegger, L., Ferreira, M., Baras, A., Reid, J., Abecasis, G., Maxwell, E., & Marchini, J. (2021). Computationally efficient whole-genome regression for quantitative and binary traits. Nature Genetics, 53(7), 1097–1103. 10.1038/s41588-021-00870-7

Mewes, H. W., Heumann, K., Kaps, A., Mayer, K., Pfeiffer, F., Stocker, S., & Frishman, D. (1999). MIPS: a database for genomes and protein sequences. In Nucleic Acids Research (Vol. 27, Issue 1). http://ziggy.sanbi.ac.za/stack/abstract.html

Nguyen, D. V, & Rocke, D. M. (2002). Tumor classification by partial least squares using microarray gene expression data. In BIOINFORMATICS (Vol. 18, Issue 1).

Ortiz, R., Reslow, F., Montesinos-López, A., Huicho, J., Pérez-Rodríguez, P., Montesinos-López, O. A., & Crossa, J. (2023). Partial least squares enhance multi-trait genomic prediction of potato cultivars in new environments. Scientific Reports, 13(1). 10.1038/s41598-023-37169-y

Ou, A. H., Rosenthal, S. B., Adli, M., Akiyama, K., Akula, N., Alda, M., Amare, A. T., Ardau, R., Arias, B., Aubry, J. M., Backlund, L., Bauer, M., Baune, B. T., Bellivier, F., Benabarre, A., Bengesser, S., Bhattacharjee, A. K., Biernacka, J. M., Cervantes, P., … Kelsoe, J. R. (2024). Lithium response in bipolar disorder is associated with focal adhesion and PI3K-Akt networks: a multi-omics replication study. Translational Psychiatry, 14(1). 10.1038/s41398-024-02811-4

Oughtred, R., Rust, J., Chang, C., Breitkreutz, B. J., Stark, C., Willems, A., Boucher, L., Leung, G., Kolas, N., Zhang, F., Dolma, S., Coulombe-Huntington, J., Chatr-aryamontri, A., Dolinski, K., & Tyers, M. (2021). The BioGRID database: A comprehensive biomedical resource of curated protein, genetic, and chemical interactions. Protein Science, 30(1), 187–200. 10.1002/pro.3978

Pers, T. H., Karjalainen, J. M., Chan, Y., Westra, H. J., Wood, A. R., Yang, J., Lui, J. C., Vedantam, S., Gustafsson, S., Esko, T., Frayling, T., Speliotes, E. K., Boehnke, M., Raychaudhuri, S., Fehrmann, R. S. N., Hirschhorn, J. N., & Franke, L. (2015). Biological interpretation of genome-wide association studies using predicted gene functions. Nature Communications, 6. 10.1038/ncomms6890

Piccininni, M., Chung, J., & Zou, L. (2022). Causal discovery in high-dimensional, multicollinear datasets.

Plenge, R. M., Scolnick, E. M., & Altshuler, D. (2013). Validating therapeutic targets through human genetics. In Nature Reviews Drug Discovery (Vol. 12, Issue 8, pp. 581–594). 10.1038/nrd4051

Qian, Y., Wang, D., Ding, Q. X., Greenberg, M., & Long, Q. (2023). The Bias of Using Cross-Validation in Genomic Predictions and Its Correction. 10.1101/2023.10.03.560782

Rabinowicz, A., & Rosset, S. (2022). Cross-Validation for Correlated Data. Journal of the American Statistical Association, 117(538), 718–731. 10.1080/01621459.2020.1801451

Rauluseviciute, I., Riudavets-Puig, R., Blanc-Mathieu, R., Castro-Mondragon, J. A., Ferenc, K., Kumar, V., Lemma, R. B., Lucas, J., Chèneby, J., Baranasic, D., Khan, A., Fornes, O., Gundersen, S., Johansen, M., Hovig, E., Lenhard, B., Sandelin, A., Wasserman, W. W., Parcy, F., & Mathelier, A. (2024). JASPAR 2024: 20thãnniversary of the open-access database of transcription factor binding profiles. Nucleic Acids Research, 52(D1), D174–D182. 10.1093/nar/gkad1059

Rodríguez-Antona, C., & Taron, M. (2015). Pharmacogenomic biomarkers for personalized cancer treatment. Journal of Internal Medicine, 277(2), 201–217. 10.1111/joim.12321

Sanseau, P., Agarwal, P., Barnes, M. R., Pastinen, T., Richards, J. B., Cardon, L. R., & Mooser, V. (2012). Use of genome-wide association studies for drug repositioning. In Nature Biotechnology (Vol. 30, Issue 4, pp. 317–320). 10.1038/nbt.2151

Solovieff, N., Cotsapas, C., Lee, P. H., Purcell, S. M., & Smoller, J. W. (2013). Pleiotropy in complex traits: Challenges and strategies. In Nature Reviews Genetics (Vol. 14, Issue 7, pp. 483–495). 10.1038/nrg3461

Strachan, T., & Read, A. (2010). Human Molecular Genetics 4th Edition. Garland Science.

Stuart, T., Butler, A., Hoffman, P., Hafemeister, C., Papalexi, E., Mauck, W. M., Hao, Y., Stoeckius, M., Smibert, P., & Satija, R. (2019). Comprehensive Integration of Single-Cell Data. Cell, 177(7), 1888–1902.e21. 10.1016/j.cell.2019.05.031

Subramanian, A., Tamayo, P., Mootha, V. K., Mukherjee, S., Ebert, B. L., Gillette, M. A., Paulovich, A., Pomeroy, S. L., Golub, T. R., Lander, E. S., & Mesirov, J. P. (2005). Gene set enrichment analysis: A knowledge-based approach for interpreting genome-wide expression profiles. www.pnas.orgcgi 10.1073/pnas.0506580102

Szklarczyk, D., Kirsch, R., Koutrouli, M., Nastou, K., Mehryary, F., Hachilif, R., Gable, A. L., Fang, T., Doncheva, N. T., Pyysalo, S., Bork, P., Jensen, L. J., & Von Mering, C. (2023). The STRING database in 2023: protein-protein association networks and functional enrichment analyses for any sequenced genome of interest. Nucleic Acids Research, 51(1 D), D638–D646. 10.1093/nar/gkac1000

Vanunu, O., Magger, O., Ruppin, E., Shlomi, T., & Sharan, R. (2010). Associating genes and protein complexes with disease via network propagation. PLoS Computational Biology, 6(1). 10.1371/journal.pcbi.1000641

Visscher, P. M., Brown, M. A., McCarthy, M. I., & Yang, J. (2012a). Five years of GWAS discovery. In American Journal of Human Genetics (Vol. 90, Issue 1, pp. 7–24). 10.1016/j.ajhg.2011.11.029

Visscher, P. M., Wray, N. R., Zhang, Q., Sklar, P., McCarthy, M. I., Brown, M. A., & Yang, J. (2017). 10 Years of GWAS Discovery: Biology, Function, and Translation. In American Journal of Human Genetics (Vol. 101, Issue 1, pp. 5–22). Cell Press. 10.1016/j.ajhg.2017.06.005

Wang, G., Sarkar, A., Carbonetto, P., & Stephens, M. (2020). A simple new approach to variable selection in regression, with application to genetic fine mapping. In J. R. Statist. Soc. B (Vol. 82). https://academic.oup.com/jrsssb/article/82/5/1273/7056114

Weeks, E. M., Ulirsch, J. C., Cheng, N. Y., Trippe, B. L., Fine, R. S., Miao, J., Patwardhan, T. A., Kanai, M., Nasser, J., Fulco, C. P., Tashman, K. C., Aguet, F., Li, T., Ordovas-Montanes, J., Smillie, C. S., Biton, M., Shalek, A. K., Ananthakrishnan, A. N., Xavier, R. J., … Finucane, H. K. (2023). Leveraging polygenic enrichments of gene features to predict genes underlying complex traits and diseases. Nature Genetics, 55(8), 1267–1276. 10.1038/s41588-023-01443-6

Wen, X., Pique-Regi, R., & Luca, F. (2017). Integrating molecular QTL data into genome-wide genetic association analysis: Probabilistic assessment of enrichment and colocalization. PLoS Genetics, 13(3). 10.1371/journal.pgen.1006646

Wilkinson, M. D., Dumontier, M., Aalbersberg, Ij. J., Appleton, G., Axton, M., Baak, A., Blomberg, N., Boiten, J. W., da Silva Santos, L. B., Bourne, P. E., Bouwman, J., Brookes, A. J., Clark, T., Crosas, M., Dillo, I., Dumon, O., Edmunds, S., Evelo, C. T., Finkers, R., … Mons, B. (2016). Comment: The FAIR Guiding Principles for scientific data management and stewardship. Scientific Data, 3. 10.1038/sdata.2016.18

Wold, H. (1975). Soft Modelling by Latent Variables: The Non-Linear Iterative Partial Least Squares (NIPALS) Approach. Journal of Applied Probability, 12(S1), 117–142. 10.1017/S0021900200047604

Wood, A. R., Esko, T., Yang, J., Vedantam, S., Pers, T. H., Gustafsson, S., Chu, A. Y., Estrada, K., Luan, J., Kutalik, Z., Amin, N., Buchkovich, M. L., Croteau-Chonka, D. C., Day, F. R., Duan, Y., Fall, T., Fehrmann, R., Ferreira, T., Jackson, A. U., … Frayling, T. M. (2014a). Defining the role of common variation in the genomic and biological architecture of adult human height. Nature Genetics, 46(11), 1173–1186. 10.1038/ng.3097

Wu, C., Zhou, F., Ren, J., Li, X., Jiang, Y., & Ma, S. (2019). A selective review of multi-level omics data integration using variable selection. In High-Throughput (Vol. 8, Issue 1). MDPI AG. 10.3390/ht8010004

Zhu, Z., Zhang, F., Hu, H., Bakshi, A., Robinson, M. R., Powell, J. E., Montgomery, G. W., Goddard, M. E., Wray, N. R., Visscher, P. M., & Yang, J. (2016). Integration of summary data from GWAS and eQTL studies predicts complex trait gene targets. Nature Genetics, 48(5), 481–487. 10.1038/ng.3538

